# DORGE: Discovery of Oncogenes and Tumor SuppressoR Genes Using Genetic and Epigenetic Features

**DOI:** 10.1101/2020.07.21.213702

**Authors:** Jie Lyu, Jingyi Jessica Li, Jianzhong Su, Fanglue Peng, Yiling Chen, Xinzhou Ge, Wei Li

**Affiliations:** Division of Computational Biomedicine, Department of Biological Chemistry, School of Medicine, University of California, Irvine, CA 92697, USA; Department of Statistics, University of California, Los Angeles, CA 90095, USA; Department of Molecular and Cellular Biology, Baylor College of Medicine, Houston, TX 77030, USA

## Abstract

Comprehensive data-driven discovery of cancer driver genes, including tumor suppressor genes (TSGs) and oncogenes (OGs), is imperative for cancer prevention, diagnosis, and treatment. Although epigenetic alterations are important contributors to tumor initiation and progression, most known driver genes were identified based on genetic alterations alone, and it remains unclear to what the extent epigenetic features would facilitate the identification and characterization of cancer driver genes. Here we developed a prediction algorithm DORGE (Discovery of Oncogenes and tumor suppressoR genes using Genetic and Epigenetic features), which integrates the most comprehensive collection of tumor genetic and epigenetic data to identify TSGs and OGs, particularly those with rare mutations. DORGE identified histone modifications as strong predictors for TSGs, and it found missense mutations, super enhancer percentages, and methylation differences between cancer and normal samples as strong predictors for OGs. We extensively validated novel cancer driver genes predicted by DORGE using independent functional genomics data. We also found that the dual-functional genes, which are both TSGs and OGs predicted by DORGE, are enriched at hubs in protein-protein interaction and drug-gene networks. Overall, our study has deepened the understanding of epigenetic mechanisms in tumorigenesis and revealed a previously undetected repertoire of cancer driver genes.

## Introduction

Cancer results from an accumulation of key genetic alterations that disrupt the balance between cell division and apoptosis (*1*). Genes with “driver” mutations that affect cancer progression are known as cancer driver genes (*2*), which can be classified as tumor suppressor genes (TSGs) and oncogenes (OGs) based on their roles in cancer progression (*3*). OGs are usually activated by gain-of-function (GoF) mutations that stimulate cell growth and division, whereas TSGs are inactivated by loss-of-function (LoF) mutations (frameshift insertions/deletions and nonsense mutations) that block TSG functions in inhibiting cell proliferation, promoting DNA repair, and activating cell cycle checkpoints.

CRISPR (Clustered Regularly Interspaced Short Palindromic Repeats)-Cas9 screens with libraries of single-guide RNAs (sgRNAs) are powerful tools for identifying genes essential for cancer cell fitness, such as cancer cell growth and viability. For example, recent CRISPR screens by the Wellcome Sanger Institute detected 628 priority targets in 324 human cell lines from 30 cancer types (*4*). However, the genes identified by CRISPR screens in cell lines, which differ vastly from primary cells, may not be physiologically relevant to human biology and disease. Indeed, many well-known cancer driver genes in the Cancer Gene Census (CGC) database (*5*) were missing in CRISPR-screening results. They might have phenotypic effects in animal models that are not included in the current CRISPR screens.

Hence, it is necessary to predict cancer driver genes based on patient genomics data. Cancer genome sequencing efforts, such as the Cancer Genome Atlas (TCGA) (*6*), have generated an unprecedentedly large data resource and enabled the development of bioinformatics algorithms to discover cancer driver genes. Tokheim *et al*. (*7*) reviewed eight major algorithms, and Bailey *et al*. (*8*) integrated 26 computational tools in a pan-cancer mutation study. These algorithms mainly look for cancer driver genes with greater than expected background mutational rates, and they output a ranked list of candidate genes based on a small collection of genetic features such as somatic mutations and copy number alterations (CNAs). Notably, Tumor Suppressor and Oncogene Explorer (TUSON) (*9*) and the 20/20+ machine-learning method (*7*) are the two major algorithms that can distinguish between protein-coding TSGs and OGs based on distinct patterns of mutational signatures.

However, a recent meta-analysis indicated that, over the next ten years, even if all available tumor genomes were analyzed, many cancer driver genes would remain undetected due to the lack of distinction between driver mutations and background mutational load (*10*). In addition, emerging evidence suggests that genetic alterations alone are insufficient to explain all cancer driver genes, including some well-known ones. For example, sustained expression of estrogen receptor-α (*ESR1*) drives two-thirds of breast cancers, but *ESR1* mutations that alter transcription levels occur in only 7% of ESR1-positive tumors (*11*). Furthermore, many pediatric tumors have extremely low mutation rates; some even appear to have no significant recurrent somatic mutations (*12*). Thus, it is likely that other mechanisms, such as epigenetic alterations, are responsible for the dysregulation of a large subset of cancer driver genes.

For example, tri-methylation on histone H3 lysine 4 (H3K4me3) and DNA methylation are the most extensively studied epigenetic modifications that influence gene expression and cell fate. H3K4me3 is a widely-recognized marker of active promoters and regulates the pre-initiation-complex formation and gene activation (*13*). More than 80% of promoters containing H3K4me3 are transcribed (*14*), and H3K4me3 is also involved in pre-mRNA splicing, recombination, DNA repair, and enhancer function. DNA methylation occurs in 70–80% of CpGs in a normal genome (*15*). H3K4me3 and CpG methylation alteration are associated with disease initiation, including many types of cancer (*16*). In particular, promoter hypermethylation that silences TSGs is a key epigenetic event in tumorigenesis (*17*), whereas gene-body methylation is positively correlated with gene expression (*18*). Recently, the “broad epigenetic domain” has emerged as a new concept in the control of cancer development. In an integrative analysis of 1,134 genome-wide ChIP-seq datasets (*19*) from the Encyclopedia of DNA elements (ENCODE) project (*20*), we found that broad H3K4me3 is a unique epigenetic signature of TSGs. In contrast to the common sharp (e.g., <1-kb width) H3K4me3 peaks associated with increased transcriptional initiation, broad H3K4me3 peaks are associated with increased transcriptional elongation. In addition, we also found many wide gene-body regions that are lowly methylated in normal tissues (the regions called “gene-body methylation canyons”) as hypermethylated in cancer (*21*). Gene-body methylation canyons are surprisingly enriched in OGs, and their hypermethylation directly induces OG activation (*21*).

Nevertheless, to the best of our knowledge, none of the existing bioinformatics algorithms sufficiently leveraged epigenetic features to predict cancer driver genes, despite the fact that epigenetic alterations are known to be associated with cancer driver genes. Therefore, these algorithms were not fully empowered, and there is a pressing need for a computational algorithm that integrates epigenetic data with genetic alterations to improve the prediction of cancer driver genes.

To address this need, we developed DORGE (Discovery of Oncogenes and tumor suppressoR genes using Genetic and Epigenetic features). DORGE includes two prediction algorithms: DORGE-TSG for predicting TSGs and DORGE-OG for predicting OGs; both algorithms are elastic-net-based logistic regression classifiers trained on CGC genes and neutral genes. By evaluating DORGE-TSG and DORGE-OG, we found a surprisingly large contribution of histone modification to TSG prediction, as well as crucial roles of the features such as missense mutations, genomics, super enhancer percentages, and hypermethylation in predicting OGs. Cancer driver genes predicted by DORGE include known cancer driver genes and novel ones that have not been reported in the literature. We evaluated these novel cancer driver genes using multiple genomics and functional genomics datasets. In addition, we found that the novel dual-functional genes, which DORGE predicted as both TSGs and OGs, are highly enriched at hubs in protein-protein interaction (PPI) and drug/compound-gene networks.

## Results

### DORGE predicts TSGs and OGs based on known cancer driver genes and neutral genes

We developed a computational tool DORGE, by integrating extensive genomic and epigenomic datasets, for predicting cancer driver genes, i.e., TSGs and OGs. Briefly, we used CGC genes and 75 curated candidate features to train two binary classification algorithms: DORGE-TSG and DORGE-OG, which we subsequently applied to every gene to predict its probability of being a TSG and OG, respectively. Finally, we used the predicted probabilities to rank genes genome-wide and identified the top-ranked genes as candidate TSGs and OGs.

Prediction of cancer driver genes is a classification problem. It requires a high-quality training dataset that contains reliable TSGs, OGs, and the genes unlikely to be TSGs or OGs. Our two positive-training gene sets include 242 TSGs and 240 OGs (with dual-functional genes removed) from the CGC database v.87., which we refer to as CGC-TSGs and CGC-OGs hereafter. The negative-training gene set includes 4,058 neutral genes (NGs) reported to have no cancer relevance (*9*). To allow for the prediction of dual-functional genes that are both a TSG and an OG, we trained two classifiers for predicting TSGs and OGs, respectively.

To develop DORGE, we constructed 75 features that are likely predictive of cancer driver genes based on the literature. These features have either known roles in TSG/OG disruption (e.g., DNA methylation, somatic mutations) or potential links to TSG/OG functions (e.g., CRISPR-screening data) (Data file S1). We categorized these features into four major types: (I) 33 mutational features from two well-known cancer driver gene prediction algorithms—TUSON (*9*) and 20/20+ (*7*)—and gnomAD; 28 out of these 33 features were compiled by TCGA (*6*) and Catalogue Of Somatic Mutations in Cancer (COSMIC) (*5*) from the mutation data of patient samples; (II) 12 genomic features including three from 20/20+ (*7*) and nine features (e.g., gene lengths and genome-evolution-related features) that have not been previously used to predict cancer driver genes (*22*); (III) 27 epigenetic features, including histone modifications from the ENCODE project (*20*), promoter and gene-body methylation features from the COSMIC database, and super enhancer percentages from the dbSUPER database (*23*); (IV) three phenotypic features, including CRISPR-screening data from the DepMap project (*24*), Variant Effect Scoring Tool (VEST) scores from 20/20+ (*7*), and gene expression Z-scores from TCGA.

To train classifiers for TSG and OG prediction, we compared eight classification algorithms: logistic regression (LR), LR with the lasso penalty, LR with the ridge penalty, LR with the elastic net penalty, random forests, support vector machines (SVM) with the linear kernel, SVM with the Gaussian kernel, and XGBoost (https://github.com/dmlc/xgboost). For each algorithm, we considered three class ratios (where a class ratio was defined as the number of NGs to the number of CGC-TSGs or CGC-OGs): the original ratio, 5:1, and 1:1; for the latter two ratios, we randomly divided NGs into partitions so that the number of NGs in each partition approximately met the ratio given the number of CGC-TSGs or CGC-OGs. Considering the imbalance between NGs and CGC-TSGs/CGC-OGs in sizes, we used the 5-fold cross validated (CV) area under the precision-recall curve (AUPRC), instead of the receiver operating characteristic curve, as the accuracy measure to compare these eight classification algorithms under the three class ratios. Our comparison result showed that downsampling the NGs to have more balanced class ratios as 5:1 and 1:1 did not improve the accuracy achieved by the original class ratio. Hence, we decided to keep the original class ratio and found that LR with the lasso, LR with the ridge, LR with the elastic net, and random forests performed the best with similar AUPRC values (Data file S2). We chose LR with the elastic net as the classification algorithm for its good interpretability and its capacity for selecting correlated, informative features. Then we trained LR with the elastic net separately for TSG and OG prediction and subsequently used the two trained algorithms to assign every gene a TSG-score and an OG-score, both ranging from 0 to 1, with a larger value indicating a higher chance of the corresponding gene being a TSG or an OG. To decide appropriate thresholds on the TSG-scores and OG-scores for final predictions, we weighted the severity of mispredicting NGs as TSGs/OGs (i.e., making false positive predictions) versus the other way around and set a target false positive rate (FPR) of 1%. Finally, we used the Neyman-Pearson classification algorithm (*25*) to set thresholds on the TSG-scores and OG-scores by respecting our target FPR and obtained two classifiers: DORGE-TSG and DORGE-OG for predicting TSGs and OGs, respectively.

Next we identified the important features for TSG and OG prediction. Because many features are correlated (Data file S1), the feature coefficients estimated by LR with the elastic net are not biologically interpretable measures of feature importance. The reason is that if one adds to the training data a feature that is highly correlated with an existing feature, the estimated coefficient of the existing feature would become less significant. This phenomenon contradicts our biological interpretation of feature importance: if a feature is important, its importance should not be diluted by the addition of another feature. Yet we are still interested in the importance of features in our final multi-feature linear classifier, so marginal feature importance based on each feature alone does not suffice. To address this issue, we proposed a simple two-step procedure: (1) we clustered features into feature groups that were approximately uncorrelated with one another; (2) we evaluated the importance of each feature group by the reduction in the 5-fold CV AUPRC when that feature group was left out, i.e., the contribution of that feature group to the 5-fold CV AUPRC given all the other feature groups. Our simple but innovative approach is advantageous in three aspects. First, by grouping correlated features we can interpret a small number of feature groups, each of which has a distinct biological interpretation, instead of a large number of features. Second, making feature groups approximately uncorrelated has a desirable consequence: if a new feature were added, it would either be added to an existing feature group or create a new feature group by itself (if it is approximately uncorrelated with any existing features); then its addition would barely affect the importance of the feature groups it is not in, as uncorrelated features would not affect each other’s importance in a multi-feature linear classifier. Third, the same criterion, 5-fold CV AUPRC, was used to select a classification algorithm and define the importance of a feature group, making the analysis self-consistent. Using this approach, we first divided all 75 features into 20 feature groups by hierarchical clustering with complete linkage so that features within each group have pairwise absolute Pearson correlations at least 0.1 (Data file S2). Then we ranked the 20 feature groups by their contributions to 5-fold CV AUPRC and selected the top-ranked groups as those whose contributions exceeded 0.005. This gave us three and five feature groups for predicting TSGs and OGs, respectively.

Analyzing these top predictive feature groups, we found that multiple histone modification features stood out as the most predictive group (whose contribution to 5-fold CV AUPRC was almost 10-fold of that of the second most predictive group containing phenotype features) for TSGs, and that missense mutations constituted the top feature group for predicting OGs (Fig. 1A and B). Besides, epigenetic features including super enhancer and promoter and gene-body cancer–normal methylation differences were among the top feature groups for predicting OGs (Fig. 1B). We also found histone modifications and missense mutations among the top predictive features for both TSGs and OGs (Fig. 1A and B), suggesting that TSGs and OGs share certain features, whose predictive power for TSGs and OGs may be different though. For each feature within a top-ranked TSG (or OG) feature group, we compared its values in the CGC-TSGs (or CGC-OGs) and the NGs by the two-sided Wilcoxon rank-sum test, and the resulting −log_10_*P*-value was shown in Fig. 1A and B.

**Fig. 1.**
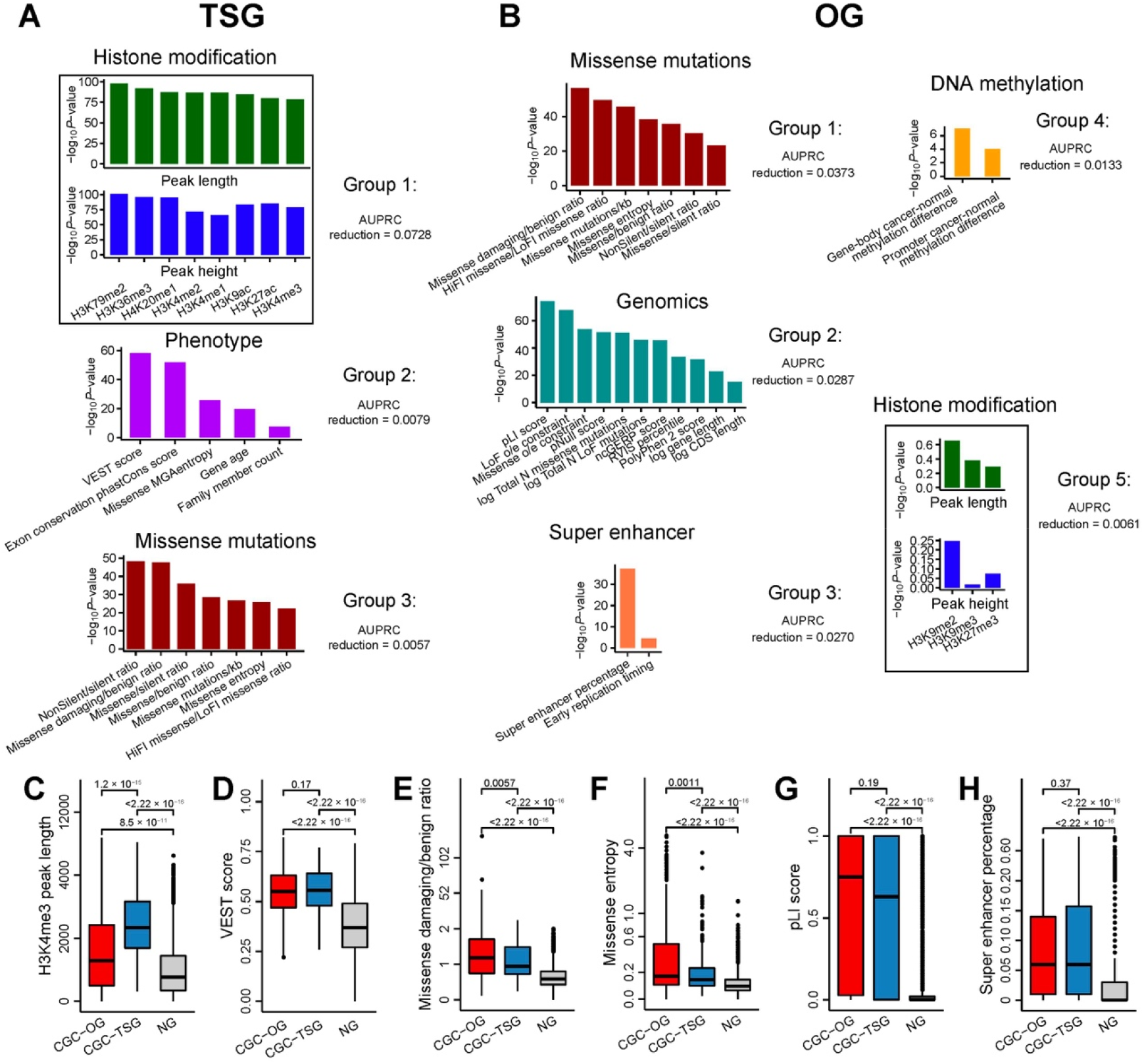
Features that discriminate tumor suppressor genes (TSGs) from oncogenes (OGs). (**A**), Feature groups selected for TSGs. (**B**), Features groups selected for OGs. Feature groups are sorted according to the AUPRC reduction in elastic net five-fold cross-validation. Feature groups are named according to the representative features. Box plots showing the distribution of (**C**), Tri-methylation on histone H3 lysine 4 (H3K4me3) mean peak length, (**D**), Variant Effect Scoring Tool (VEST) score, (**E**), Missense damaging/benign ratio, (**F**), Missense entropy, (**G**) pLI score and (**H**), Super enhancer percentage for the CGC-OG, CGC-TSG, and NG sets. Genes as both TSGs and OGs are excluded. *P*-values for the differences between the TSGs/OGs and NGs were calculated by the one-sided “greater-than” Wilcoxon rank-sum test.

We further examined several features in terms of their individual, marginal power of distinguishing CGC-TSGs and CGC-OGs from NGs. Indeed, multiple features are marginally strong predictors of TSGs, as they have significantly higher values in CGC-TSGs than in NGs. They include epigenetic features such as H3K4me3 peak length and height (Fig. 1C and Fig. S1A) and H3K79me2 peak length and height (Fig. S1B and S1C), missense mutational features such as non-silent/silent ratio (Fig. S1D), and phenotype features such as Variant Effect Scoring Tool (VEST) score (Fig. 1D). Many features also have significantly higher values in CGC-OGs than in NGs. They include missense mutational features such as missense damaging/benign ratio (Fig. 1E), missense entropy (Fig. 1F), probability of being loss-of-function intolerant (pLI) score (Fig. 1G) and LoF o/e constraint (Fig. S1E), genomics features such as evolutionary conservation phastCons score and non-coding Genomic Evolutionary Rate Profiling (ncGERP) score (Fig. S1F and S1G), and epigenetic features such as super enhancer percentage in cell-lines (Fig. 1H). In particular, our finding agrees with previous studies in that missense damaging/benign ratio (reflecting the functional impact of missense mutations) and missense entropy (representing the enrichment of mutations in few residues) (*9*) have significantly higher values in CGC-OGs than in CGC-TSGs and NGs (Fig. 1E and F). Interestingly, VEST and PolyPhen-2 scores, both of which reflect functional effects of mutations, have significantly higher values in CGC-TSGs and CGC-OGs than in NGs, and they do not exhibit statistically significant differences between CGC-TSGs and CGC-OGs (Fig. 1D and S1H). Notably, we found super enhancer, a commonly regarded OG-specific feature (*26*), also characteristic of TSGs, as it has significantly higher values in CGC-TSGs than in NGs (Fig. 1H).

We note that, besides H3K4me3 peak length, a readily known TSG predictor, peak lengths of four more histone marks (H3K79me2, H3K36me3, H4K20me1, and H3K9ac) are also significantly larger in CGC-TSGs than in CGC-OGs and NGs (Fig. S1B, S1I, S1J, and S1K), consistent with the fact that the activation of TSGs is associated with transcriptional elongation (*19, 27–29*). To further verify the enrichment of broad H3K4me3 peaks in CGC-TSGs, we performed the Fisher’s exact test on a two-by-two contingency table, whose two rows correspond to CGC-TSGs and all the other genes in the training data (CGC-OGs and NGs) and whose two columns correspond to the genes with broad H3K4me3 peaks (whose mean lengths across ENCODE samples > 4 kb) and the rest of genes. We similarly performed two more Fisher’s exact tests to check the enrichment of broad H3K4me3 peaks in CGC-OGs and NGs but found much lower enrichment in these two gene groups than in CGC-TSGs, confirming that H3K4me3 is a distinctive feature of TSGs (Fig. S1L). Taken together, we identified histone modifications as the top predictors for TSGs. We found missense mutations, super enhancer percentages, and methylation differences between cancer and normal samples as major predictors for OGs. It is worth noting that histone modifications and missense mutations are also important features for predicting OGs and TSGs, respectively, though to a lesser extent. In summary, DORGE can successfully leverage public data to discover the genetic and epigenetic alterations that play significant roles in cancer driver gene dysregulation. Fig. S2 provides an overview of the DORGE method and the evaluations in the following sections.

### Evaluation of the prediction accuracy of DORGE

As we described earlier, DORGE-TSG and DORGE-OG output TSG-scores and OG-scores for predicting TSGs and OGs, respectively. Every gene received a TSG-score and an OG-score, both ranging from 0 to 1, and a higher TSG-score (or OG-score) indicates a higher probability of a gene being a TSG (or an OG) (Materials and Methods). DORGE thresholded the TSG-scores and OG-scores by the Neyman-Pearson classification algorithm (*25*) with a target FPR of 1%, leading to 925 predicted TSGs, whose TSG-scores exceeded 0.6233374, and 683 predicted OGs, whose OG-scores exceeded 0.6761319. In total, DORGE predicted 1,172 cancer driver genes, including 436 dual-functional genes (Fig. 2A; the predicted genes are listed in Data file S2). We note that these predicted TSGs and OGs are conservative predictions guided by the small FPR threshold 1%, as reflected by the fact that their numbers are smaller than the numbers of previously predicted cancer driver genes—1,217 TSGs and 803 OGs in databases TSGene (*30*) and ONGene (*31*) (by June 18, 2020). If DORGE users would like to be less conservative and predict more TSGs and OGs, they can opt for a higher FPR threshold such as 5%. Next, we filtered out CGC genes from the DORGE-predicted cancer driver genes and defined the remaining 725 predicted TSGs and 515 predicted OGs as DORGE-predicted novel genes (Data file S1), among which 537 novel TSGs were not included in the CancerMine (*32*) or TSGene database (Fig. 2B), and 306 novel OGs were not found in the CancerMine or ONGene database (Fig. 2C).

**Fig. 2.**
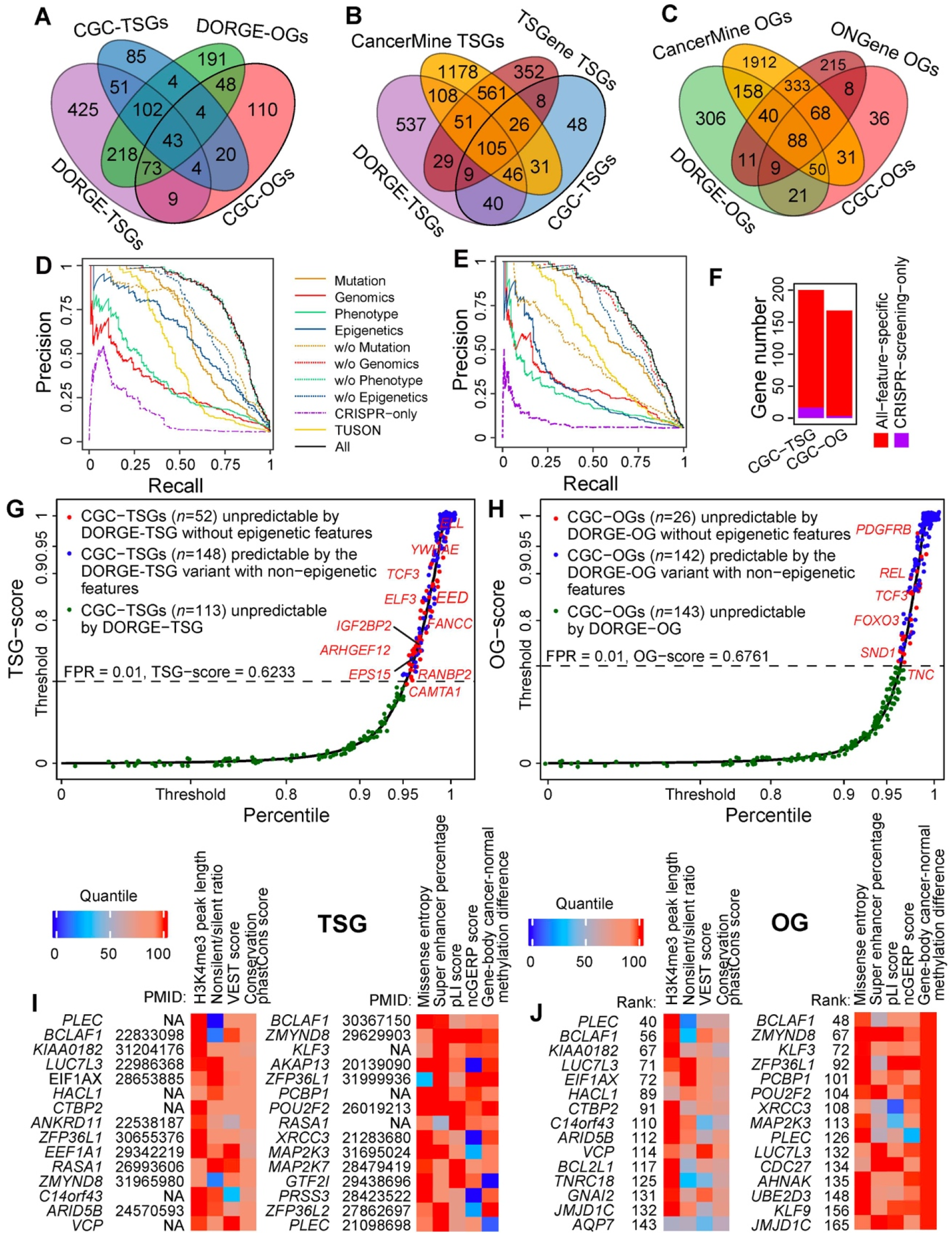
Evaluation of the DORGE method and characterization of the DORGE-predicted novel TSGs and OGs. Venn diagrams showing the overlap (**A**), between DORGE-predicted novel TSGs/OGs and CGC-TSGs/OGs. (**B**), between DORGE-predicted novel TSGs, CGC-TSGs, CancerMine-TSGs, and TSGene database-TSGs. (**C**), between DORGE-predicted novel OGs, CGC-OGs, CancerMine-OGs, and ONGene database-OGs. Precision-recall curves (PRCs) for (**D**), TSG and (**E**), OG prediction. Different lines represent different PRCs from DORGE or DORGE variants. (**F**), Stacked bar plots showing the number of rediscovered CGC-TSGs and CGC-OGs using all features compared to CRISPR-screening data only. Cumulative distribution function (CDF) plots of DORGE-predicted TSG-scores (**G**) and OG-scores (**H**) of 19,636 human genes. X-axis and Y-axis are swapped for illustration purposes, and Y-axis is stretched to emphasize large TSG- and OG-scores. CGC genes are plotted as Jitter points to avoid overplotting. The dashed lines indicate DORGE-TSG and DORGE-OG thresholds at a target FPR of 1%, and the CGC genes whose TSG-scores and OG scores exceed the thresholds (above the dashed lines) are predicted as TSGs and OGs. (**I**), Top-15 DORGE-predicted non-CGC novel TSGs (left) and OGs (right), respectively, along with representative feature heatmaps and PubMed IDs. To make features comparable, feature values are transformed into quantiles. (**J**), Top-15 DORGE-predicted non-CGC novel TSGs (left) and OGs (right) that have no documented role in cancer based on the TSGene, ONGene, and CancerMine databases, along with representative feature heatmaps.

We evaluated DORGE-TSG and DORGE-OG by their overall prediction accuracy and found that they achieved high 5-fold CV AUPRC of 0.821 and 0.766, respectively, when trained with all the 75 features (Fig. 2D and 2E). Considering that previous algorithms primarily relied on genetic features to predict cancer driver genes, we evaluated the accuracy gain of DORGE from including epigenetic and phenotypic features. To this end, we constructed variants of DORGE-TSG and DORGE-OG based on each of the following feature subsets: ‘Mutation’, ‘Genomics’, ‘Phenotype’, ‘Epigenetics’, and their complements (i.e., the subsets resulting from subtracting each of the four feature subsets from the 75 features), as well as TUSON and CRISPR-screening-only features (Data file S1, Fig. 2D and E). For each of these DORGE-TSG and DORGE-OG variants, we calculated its 5-fold CV AUPRC.

Based on feature subsets ‘Mutation’, ‘Genomics’, ‘Phenotype’, and ‘Epigenetics’, the corresponding DORGE-TSG variants achieved 5-fold CV AUPRC of 0.638, 0.314, 0.358, and 0.600, respectively. In parallel, based on the complements of ‘Mutation’, ‘Genomics’, ‘Phenotype’, and ‘Epigenetics’ (i.e., when features in each subset were excluded), the corresponding DORGE-TSG variants achieved 5-fold CV AUPRC of 0.692, 0.819, 0.820, or 0.715. These results consistently show the large contributions of ‘Mutation’ and ‘Epigenetics’ features to TSG prediction (Fig. 2D). Furthermore, using the features in the TUSON method and the CRISPR-screening-only feature, the corresponding DORGE-TSG variants only achieved 5-fold CV AUPRC of 0.500 and 0.156, much lower than 0.821 achieved by DORGE-TSG with all the 75 features. Similarly, we compared DORGE-OG with its variants trained on feature subsets. Specifically, DORGE-OG variants that only used ‘Mutation’, ‘Genomics’, ‘Phenotype’, or ‘Epigenetics’ features achieved 5-fold CV AUPRC of 0.660, 0.299, 0.241, or 0.295; when each of these feature subsets was excluded, the AUPRC correspondingly became 0.453, 0.752, 0.763, or 0.705. These results suggest that ‘Mutation’ features have a large contribution to OG prediction (Fig. 2E). Similar to DORGE-TSG, the DORGE-OG variants trained with TUSON features or the CRISPR-screening-only feature had much lower prediction accuracy (5-fold CV AUPRC of 0.534 or 0.089) than that of DORGE-OG trained with all the 75 features (5-fold CV AUPRC of 0.766). The fact that DORGE-TSG and DORGE-OG outperformed all their variants confirms that DORGE effectively leveraged the 75 features and did not suffer from overfitting in its TSG and OG prediction.

The above results also reveal that the CRISPR-screening-only feature did not have a high predictive power on its own, as shown by its low 5-fold CV AUPRC (0.156 and 0.089) in TSG and OG prediction. Moreover, under the target FPR of 1%, the DORGE-TSG and DORGE-OG variants with the CRISPR-screening-only feature identified only 16 (5.1%) CGC-TSGs and 3 (1.0%) CGC-OGs, whereas DORGE-TSG and DORGE-OG with all the 75 features recovered additional 184 (58.8%) CGC-TSGs and 165 (53.1%) CGC-OGs (Fig. 2F). These results challenge a common belief that CRISPR screening using cell lines is powerful for discovering cancer driver genes. A possible reason for our results is that cell lines do not well mimic *in vivo* cancer cells. These additional cancer driver genes with all the 75 features might have phenotypic effects in animal models that are not included in the current CRISPR screens

We next evaluated the distinct predictive power provided by epigenetic features to cancer driver gene prediction. Inspecting the distributions of TSG-scores and OG-scores, we found that many top-ranked CGC genes were not predictable by DORGE without epigenetic features (Fig. 2G and H). In detail, 52 (16.61%) CGC-TSGs and 26 (8.36%) CGC-OGs would have been missed by DORGE-TSG and DORGE-OG, respectively, at the target FPR 1% if epigenetic features were not included. These results suggest that (I) epigenetic features empowered the discovery of cancer driver genes; and (II) epigenetic features empowered DORGE-TSG more than DORGE-OG because the number of rescued CGC-TSGs (52) is twice the number of rescued CGC-OGs (26).

We then searched biomedical literature for the top-15 novel TSGs and OGs ranked by DORGE. Out of these top novel genes, 10 TSGs and 12 OGs have reported tumor suppressive and oncogenic functions, respectively (Fig. 2I). We also inspected these top novel genes for selected representative features and confirmed that they indeed have high values in the top predictive TSG features (H3K4me3 peak length, nonsilent/silent ratio, VEST score, and conservation phastCons score) and OG features (missense entropy, super enhancer percentage, pLI score, ncGERP score, and gene-body cancer–normal methylation difference) selected from the top feature groups (Fig. 2I). We further confirmed this result in the subset of top novel genes that are not in the CancerMine, TSGene, and ONGene databases (Fig. 2J). In particular, nearly all of the top novel TSGs have broad H3K4me3 peaks, and most of the top novel OGs are hypermethylated in gene-body (with positive cancer–normal methylation differences).

### Benchmarking DORGE against existing algorithms

We further compared DORGE with ten existing algorithms for cancer driver gene prediction using four accuracy measures—sensitivity (*Sn*), specificity (*Sp*), precision, and overall accuracy—all based on CGC genes (Table 1). We did not include the five-test model (RF5) because even though it outputs TSG and OG probabilities, it does not have explicit cutoffs for defining TSGs and OGs (*33*). We found that DORGE performed the best in all these measures except *Sp*, for which DORGE was 0.997 and the best algorithm 20/20+ was 1.000. The superiority of DORGE was most obvious in *Sn*, where its top performance (0.611) was followed with a large gap by OncodriveFM (0.338) (*34*), MuSIC (0.331) (*35*), and MutPanning (0.318) (*36*) (Table 1). To further confirm that DORGE outperformed these ten algorithms, we performed a similar comparison based on 1,056 OncoKB cancer genes (*37*), which had been widely used to benchmark cancer gene prediction. Consistent with the CGC gene evaluation results, DORGE achieved the best performance in *Sn* (almost 50% higher than that of the second best algorithm OncodriveFM) and overall accuracy, the third best performance in *Sp* (0.997 vs. 0.999 of the best method TUSON), and the second best performance in precision (0.973 vs. 0.993 of the best method 20/20+, whose *Sn* was only 32% of that of DORGE) (Data file S2). Taken together, our benchmark results show that DORGE made a significant advance in improving cancer driver gene prediction from existing algorithms.

**Table 1.** Evaluation of cancer driver genes (TSGs + OGs) prediction based on the v.87 CGC genes.

| Method | # | *Sn* | *Sp* | Precision | Accuracy | Algorithms |
| --- | --- | --- | --- | --- | --- | --- |
| DORGE | 1,172 | 0.611 | 0.997 | 0.966 | 0.948 | Logistic regression with the elastic net model |
| OncodriveFM (34) | 2,600 | 0.338 | 0.915 | 0.367 | 0.841 | Functional impact model |
| MuSIC (35) | 1,975 | 0.331 | 0.870 | 0.272 | 0.801 | Mutational background model |
| MutPanning (36) | 460 | 0.318 | 0.994 | 0.880 | 0.907 | Nucleotide context model |
| TUSON (9) | 243 | 0.222 | 0.999 | 0.961 | 0.900 | *P*-value combination |
| OncodriveFML (58) | 680 | 0.212 | 0.983 | 0.646 | 0.885 | Functional impact model |
| 20/20+ (7) | 193 | 0.208 | 1.000 | 0.991 | 0.899 | Random Forest model |
| GUST (78) | 276 | 0.206 | 0.994 | 0.838 | 0.894 | Random Forest model |
| MutSigCV (57) | 158 | 0.137 | 0.998 | 0.905 | 0.888 | Mutational background model |
| OncodriveCLUST (59) | 586 | 0.118 | 0.963 | 0.319 | 0.855 | Mutational hotspot model |
| ActiveDriver (61) | 417 | 0.098 | 0.996 | 0.771 | 0.881 | Logistic regression model |

Based on CGC-TSGs and CGC-OGs, we further benchmarked DORGE against 20/20+, TUSON, and GUST for separate prediction of TSGs and OGs (Data file S2). We did not include the other seven algorithms because they could not predict TSGs and OGs separately. Consistent with our previous results, DORGE exhibited much higher *Sn* than the other three algorithms did (DORGE had *Sn* of 0.639 and 0.54 for predicting TSGs and OGs, while the best *Sn* of the other three algorithms was only 0.252 and 0.116), and it also achieved the best precision and overall accuracy; all the four algorithms had close to perfect *Sp*. Although the high *Sn* of DORGE seemed to be due to the fact that 20/20+, TUSON, and GUST by default predicted fewer TSGs and OGs than DORGE did, it was not the case. After we adjusted the thresholds of 20/20+ and TUSON so that they predicted the same numbers of TSGs and OGs as DORGE did (the GUST software does not allow such threshold adjustment), the *Sn* of 20/20+ and TUSON, though increased, remained almost one-fold lower than that of DORGE. Collectively, our results suggest that DORGE outperformed 20/20+, TUSON, and GUST in both TSG and OG prediction.

We also compared DORGE with TUSON and 20/20+ in terms of their predicted ranking of CGC-TSGs and CGC-OGs. For example, if an algorithm predicted gene A more likely than gene B to be a TSG, we say that gene A received a smaller TSG rank than gene B did. Accordingly, we calculated a TSG rank and an OG rank for every CGC gene by each algorithm. Among the CGC genes, we define the core CGC-TSGs and core CGC-OGs as those that were annotated solely as TSGs and OGs, not both (dual-functional), in CGC v.77. Compared to the genes that were added later to CGC v.87, these core CGCs have been more extensively studied. Then we examined the ranking consistency between DORGE and the other two algorithms for CGC genes and the core CGC genes. For CGC-TSGs, we found that their TSG ranks by DORGE had strong positive correlations with their TSG ranks by TUSON and 20/20+ (Fig. S3A and S3B), and overall they were ranked more top by DORGE than by the other two algorithms (Fig. S3E). We observed similar results for CGC-OGs (Fig. S3C, S3D, and S3G). The conclusions also held for core CGC genes (Fig. S3F and S3H). These results confirm that DORGE predictions are more biologically relevant than those of TUSON and 20/20+. For example, *ELL* (elongation factor for RNA polymerase II), a CGC-TSG, was ranked 190-th by DORGE-TSG, 8,144-th by TUSON, and 3,958-th by 20/20+; *PDGFB* (platelet derived growth factor subunit B), a CGC-OG, was ranked 207-th by DORGE, 2,753-th by TUSON, and 4,982-th by 20/20+. Also, DORGE ranked CGC dual-functional genes better than TUSON and 20/20+ did, as exemplified by the dual-functional gene *IDH1* (isocitrate dehydrogenase (NADP(+)) 1), which was ranked first for TSG and 28-th for OG by DORGE, 18,734-th for TSG and 2,092-th for OG by TUSON, and 14,936-th for TSG and 13-th for OG by 20/20+.

### Functional evaluation of novel cancer driver genes and those unpredictable without epigenetics features

Even though DORGE predicted many more cancer driver genes than TUSON, 20/20+, and GUST did— DORGE, TUSON, 20/20+, and GUST predicted 1,172, 243, 193, and 276 cancer driver genes, respectively, DORGE achieved the highest overall prediction accuracy based on CGC genes. After confirming this, we further characterized the novel cancer driver genes, defined as those predicted by DORGE but not included in the CGC database.

We performed the Kyoto Encyclopedia of Genes and Genomes (KEGG) pathway analysis on the novel TSGs and OGs, and we found, as expected, that the novel TSGs are enriched with TSG-related pathways such as “apoptosis” and “focal adhesion” and that the novel OGs are enriched with OG-related pathways such as “cell cycle” and “TGF-beta signaling pathway” (Fig. 3A). However, without epigenetic features, the novel TSGs and OGs predicted by the DORGE-TSG and DORGE-OG variants are no longer enriched with certain TSG-related and OG-related pathways such as “TGF-beta signaling pathway” (Fig. S4A). These results again suggest that epigenetic features made unique contributions to discovering novel cancer driver genes. In addition, the degrees of enrichment (−log_10_*P*-values) of those shared enriched KEGG pathways, which were enriched in novel TSGs or OGs regardless of the inclusion of epigenetic features, are positively correlated, implying that the addition of epigenetic features did not prohibit the discovery of meaningful cancer driver genes (Fig. S4B and S4C).

**Fig. 3.**
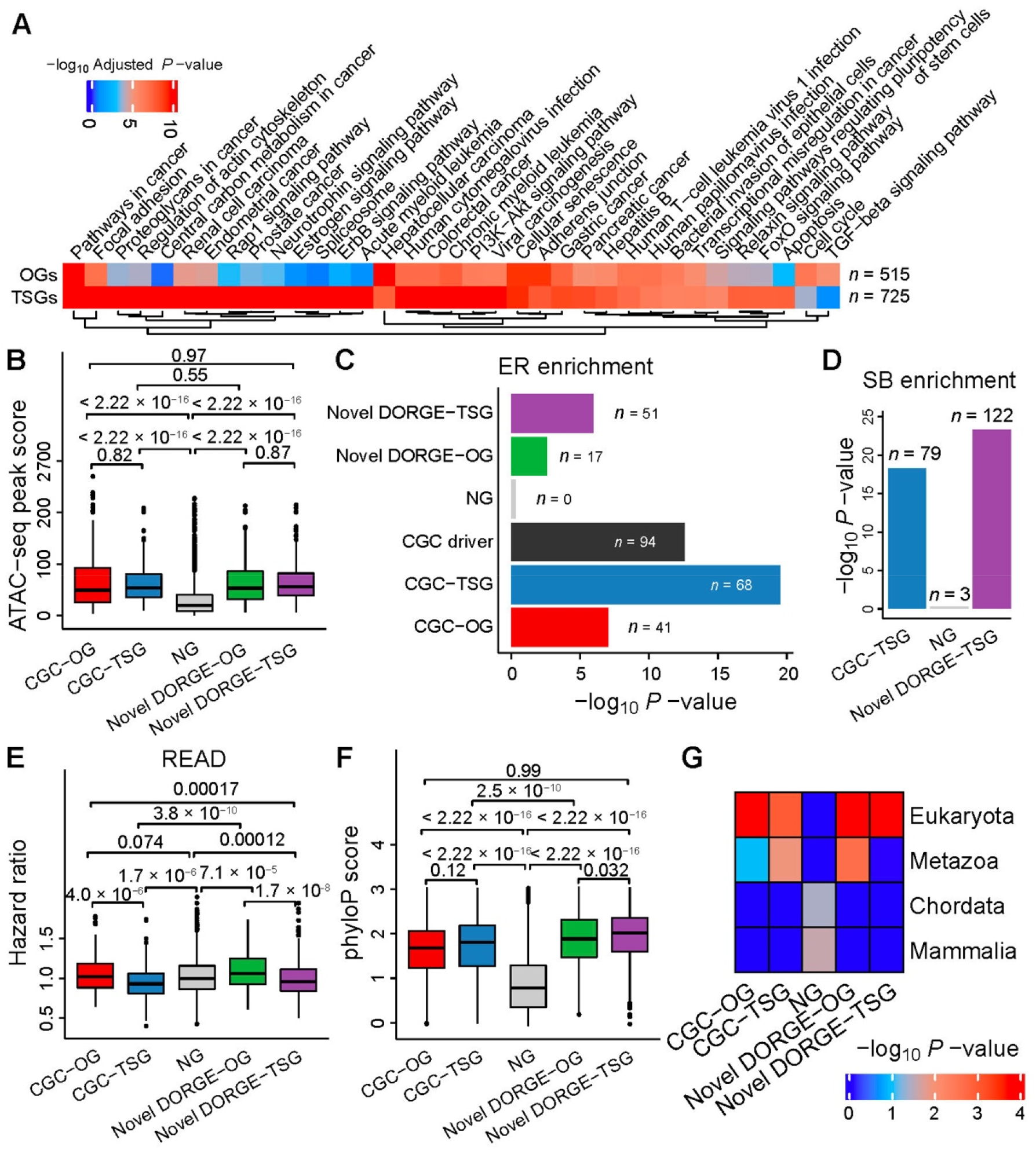
Characterization and evaluation of DORGE-predicted novel TSGs/OGs by independent functional genomic and genomic datasets. (**A**), Kyoto Encyclopedia of Genes and Genomes (KEGG) pathway enrichment analysis performed by Enrichr (*75*) for DORGE-predicted novel TSGs and OGs. Due to space limitations, terms with adjusted *P*-values < 10^−4^ are shown. Besides, terms with adjusted *P*-values 10^8^-fold lower for TSGs than OGs or 10^4^-fold lower for OGs than TSGs are also shown. (**B**), ATAC-seq peak score measuring open chromatin for CGC-TSGs/OGs, DORGE-predicted novel TSGs/OGs, and NGs. Enrichment heatmaps of various gene types in (**C**), epigenetic regulator (ER) gene list and (**D**), inactivating pattern gene list for Sleeping Beauty insertional mutagenesis, a screening tool for cancer driver genes. (**E**), Boxplot showing the Cox hazard ratio (HR) score for various gene types. Data are from Rectum adenocarcinoma (READ). (**F**), Boxplot showing the phyloP score for various gene types. The phyloP score measures phylogenetic conservation and represents −log*P*-values under a null hypothesis of neutral evolution. PhyloP basewise conservation scores were derived from a Multiz alignment of 46 vertebrate species. (**G**), TSGs and OGs are enriched in genes having earlier evolutionary origin (Eukaryota). *P*-values for the differences between indicated gene categories were calculated by the one-sided Wilcoxon rank-sum test. In boxplots and heatmap, the Fisher’s Exact Test is used to calculate *P*-values, and gene numbers in different gene categories are normalized to 200 to make *P*-values comparable. In this figure, dual-functional CGC genes were excluded from the CGC-TSGs/OGs.

Given that histone modification features (e.g., H3K4me3 peak length) empowered DORGE-TSG prediction, we sought experimental evidence for the novel TSGs that have broad histone modification (e.g., H3K4me3) peaks. A previous cell proliferation experiment observed increased cell growth after knocking down multiple potential TSGs whose H3K4me3 peaks have mean lengths (across ENCODE cell lines) greater than 2 kb (*19*), including two DORGE predicted novel TSGs—*CSRNP1* and *NR3C1*B. Another previous study found that *Mll4* loss downregulates potential TSG expression and weakens broad H3K4me3 peaks in mice (*38*). Examining the human orthologs of the six mouse potential TSGs downregulated by *Mll4* loss in that study, we found that four orthologs were ranked top by DORGE-TSG and have H3K4me3 peaks longer than 2 kb. These four human genes are *DNMT3A* (18-th), *BCL6* (96-th), *FOXO3* (222-th), and *CBFA2T3* (1,012-th).

### Characterization of DORGE-predicted novel TSGs and OGs by independent functional genomics data

We first used a published ATAC-seq dataset of TCGA pan-cancer samples (*39*) to characterize the DORGE-predicted novel cancer driver genes. ATAC-seq reveals gene accessibility and provides valuable information about the complex gene regulatory relationships. Based on this ATAC-seq dataset, we found that DORGE-predicted novel TSGs and OGs—consistent with that CGC-TSGs and CGC-OGs—are significantly more accessible than NGs (all with *P* = 2.22 × 10^−16^ by the one-sided Wilcoxon rank-sum test) (Fig. 3B). This result established a connection between cancer driver genes and chromatin accessibility— both TSGs and OGs are ubiquitously accessible in cancer samples.

We then explored a possible relationship between cancer driver genes and epigenetic regulators (ERs), which are known to play fundamental roles in genome-wide gene regulation by reading or modifying chromatin states. A previous study suggested that most ERs are intolerant to LoF mutations (*40*), and our Fig. S1E also shows that LoF mutations (reflected by the LoF o/e constraint feature) are significantly more abundant in TSGs and OGs than NGs, prompting us to explore whether ER genes have a significant overlap with cancer driver genes. By analyzing a curated list of 761 ERs, we found significant enrichment of CGC-TSGs and CGC-OGs (*P* = 3.14 × 10^−20^ and 9.36 × 10^−8^ by the Fisher’s exact test; in total, 94 CGC cancer driver genes are among the ERs, with *P* = 2.79 × 10^−13^ by the Fisher’s exact test) (Fig. 3C). This result also shows the greater enrichment of CGC-TSGs than that of CGC-OGs in ER genes, consistent with a previous study showing that the application of cancer gene classifiers to ER genes revealed more TSGs than OGs (*41*). Notably, similar to CGC genes, DORGE-predicted novel TSGs (*P* = 1.15 × 10^−6^) are also more enriched than novel OGs (*P* = 2.65 × 10^−3^) in ER genes (Fig. 3C).

We next evaluated DORGE-predicted novel TSGs using Sleeping Beauty (SB) screening data. The SB transposon is a type of synthetic DNA elements that can disrupt the expression of genes near its insertion sites, a process called insertional mutagenesis. Hence, the SB transposon is a screening tool for TSGs, whose expression disruption leads to carcinogenesis. To verify the novel TSGs, we downloaded the list of inactivating pattern genes from the Sleeping Beauty Cancer Driver Database (SBCDDB) (*42*). As expected, we found that both CGC-TSGs (*P* = 5.41 × 10^−19^) and DORGE-predicted novel TSGs (*P* = 5.11 × 10^−24^) are enriched in the list. In contrast, NGs have no enrichment. This result is consistent with our expectation that TSGs are inactivated in SB screens (Fig. 3D).

We further evaluated DORGE-predicted novel cancer driver genes using an shRNA screening dataset from the Achilles project (*43*), as shRNA screens for gene essentiality for cell proliferation in cell lines. Based on the dataset, the knockdown of DORGE-predicted novel OGs and CGC-OGs shows a greater decrease in cell proliferation rates compared to NGs (Fig. S4D). In contrast, the knockdown of DORGE-predicted novel TSGs and CGC-TSGs shows nearly no decrease in cell proliferation rates compared to NGs (Fig. S4D). This result is consistent with the prior knowledge that the proliferation of cell lines is dependent upon OGs (*24*).

Lastly, we evaluated DORGE-predicted novel cancer driver genes using patient survival data. In the precomputed survival data downloaded from the OncoRank website (*44*), every gene has a hazard ratio (HR, whose value >, =, or < 1 indicates that the gene’s expression reduces, does not affect, or increases patients’ survival time). We found that CGC-TSGs and DORGE-predicted novel TSGs have significantly lower HRs than OGs (CGC-OGs and DORGE-predicted novel OGs) and NGs in three representative cancer types: Rectum adenocarcinoma (READ), Colon adenocarcinoma (COAD), and Uterine Corpus Endometrial Carcinoma (UCEC) (Fig. 3E, S4E and S4F). These results are consistent with the fact that TSG expression prohibits cancer occurrence and prolongs survival, while OG expression has the opposite effects. The complete HRs and *P*-values of DORGE-predicted novel TSGs and OGs in 21 cancer types are available in Data file S1.

### TSGs and OGs are conserved at both exons and non-coding regions

Previous studies have suggested that evolutionarily conserved genes are enriched with cancer driver candidates and drug targets (*45*). Consistent with these studies, we observed statistically significant differences in exonic sequence conservation (phastCons and phyloP scores) between CGC-TSGs/OGs and NGs, and the same conclusion holds for DORGE-predicted TSGs and OGs (Fig. 3F and S4G). Compared to OGs, TSGs have slightly higher exonic sequence conservation (Fig. 3F and S4G).

We next explored the conservation of non-coding regions in cancer driver genes. Non-coding regions are characterized by positive non-coding Genomic Evolutionary Rate Profiling (ncGERP) values and negative non-coding Residual Variation Intolerance Score (ncRVIS) values. The reason is that ncGERP is a measure of nucleotide constraints and reflects conservation across the mammalian lineage (*46*) (Fig. S1G), while ncRVIS measures human-specific constraints (*46*). Based on these two measures, we found that TSGs (CGC-TSGs and DORGE-predicted novel TSGs) are slightly more conserved than OGs (CGC-OGs and DORGE-predicted novel OGs) at non-coding regions (Fig. S1G and Fig. S4H).

In summary, we found that cancer driver genes are more conserved than NGs at both exonic and non-coding regions. Between TSGs and OGs, we, for the first time to our knowledge, found that TSGs are more conserved at exons, while OGs are more conserved at non-coding regions.

### TSGs and OGs are overrepresented in ancient genes

Motivated by our conservation results, we investigated the phyletic ages (i.e., evolutionary origins) of cancer driver genes. Although cancer driver genes are believed to be originated from Metazoa (multicellular animals) (*47*), the possibility of their origination from Eukaryota, an earlier evolutionary origin, has not been explicitly investigated. Based on the phyletic-age gene lists (from early to late: Eukaryota, Metazoa, Chordata, and Mammalia) from the Online GEne Essentiality (OGEE) database (*48*), we found significant enrichment of cancer driver genes in the Eukaryota gene list (Fig. 3G; *P*-values by the Fisher’s exact test: *P* = 1.05 × 10^−3^ for CGC-TSGs, *P* = 3.25 × 10^−13^ for DORGE-predicted novel TSGs, *P* = 1.41 × 10^−5^ for CGC-OGs, and *P* = 2.77 × 10^−5^ for DORGE-predicted novel OGs), in contrast to NGs. Our results indicate that cancer driver genes may be originated earlier in the evolutionary history than previously thought. In addition, we found that cancer driver genes were not enriched in young phyletic ages (Chordata and Mammalia) (Fig. 3G), consistent with a recent paper (*49*).

### Dual-functional cancer driver genes act as backbones in protein-protein interaction networks

Previous studies have shown high interactivity of cancer driver genes in the BioGRID PPI network (*9*), and accordingly, PPI data have been used to identify cancer driver genes (*50, 51*). We, therefore, explored the extent to which DORGE-predicted TSGs and OGs are connected to other genes/proteins. When analyzing the whole BioGRID PPI network (Fig. 4A), we found that TSGs and OGs, including CGC genes and DORGE-predicted novel genes, exhibit significantly higher degrees, betweenness, and closeness centrality than NGs do (Fig. S5A–C). This result suggested that the removal or knockdown of cancer driver genes, as expected, will exert a critical impact on the whole PPI network. In particular, dual-functional driver genes as both TSGs and OGs display even higher interactivity than sole TSGs and OGs (Fig. S5A–C). Densely connected genes tend to form modules, and importantly, cancer driver gene modules can trigger the hallmarks of cancer and confer the proliferation advantages displayed on cancer cells (*52*). Here, we used the Molecular Complex Detection (MCODE) algorithm to identify six densely connected network modules/backbones (Fig. 4B) from the PPI subnetwork of the 1,172 DORGE-predicted cancer driver genes. Notably, the 64 genes that comprise the six identified modules are all dual-functional genes (8 CGC dual-functional genes and 56 DORGE-predicted novel dual-functional genes). This overrepresentation of dual-functional driver genes in network modules is unusual, as it is highly unlikely to obtain a 64-gene subnetwork comprised of all dual-functional genes (*P* = 6.66 × 10^−27^ by the binomial test).

**Fig. 4.**
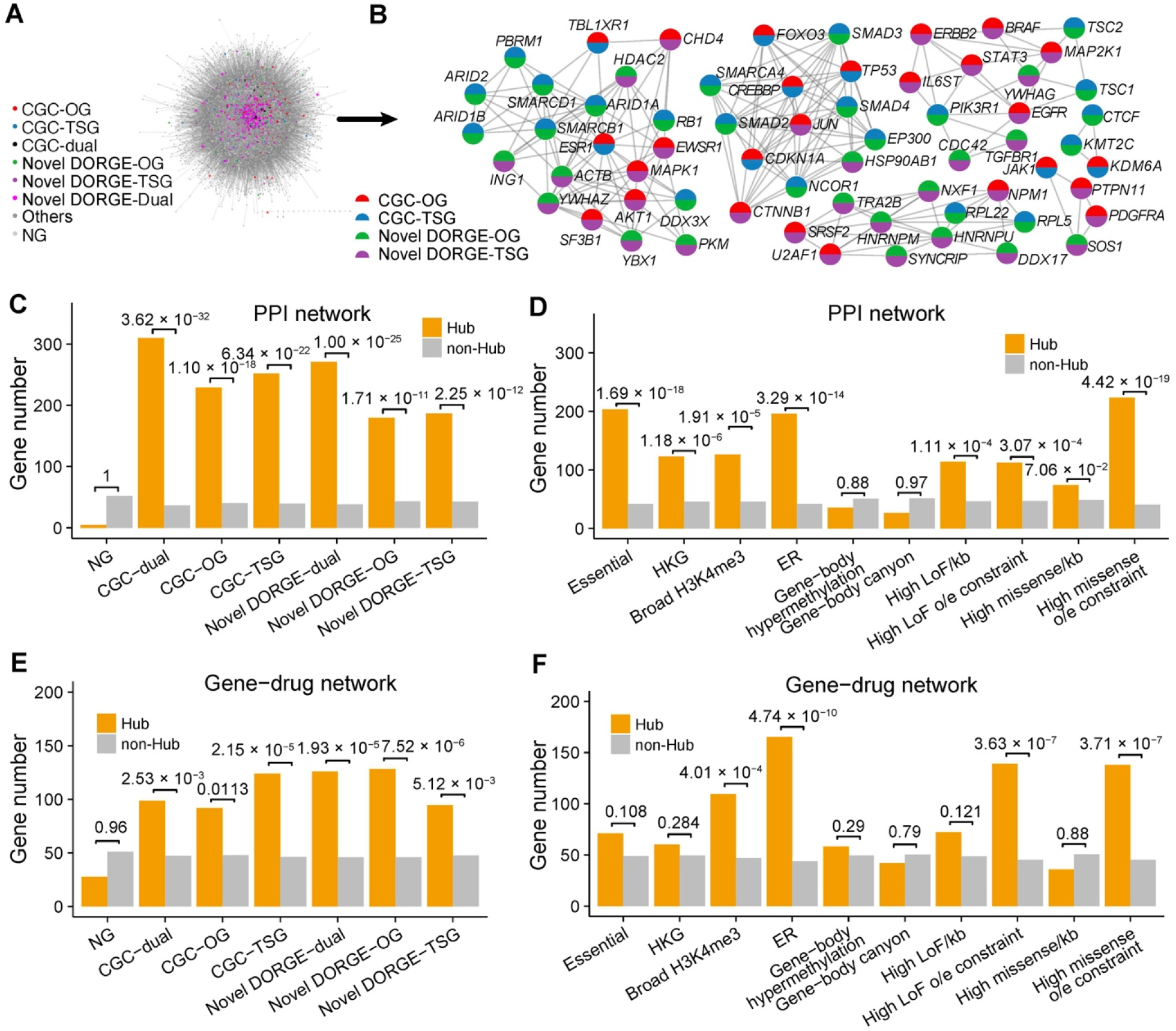
Dual-functional cancer driver genes act as backbones in BioGRID protein-protein interaction (PPI) and characterization of hub genes in PPI and PharmacoDB gene-drug networks. (**A**), Complete BioGRID PPI network. (**B**), The Molecular Complex Detection (MCODE) algorithm was applied to DORGE-predicted novel TSGs/OGs to identify densely connected network modules (or backbones). All genes in the identified network are CGC dual-functional genes or novel dual-functional genes. Gene categories are represented as pie charts, with the colors coded based on gene categories. (**C**), Enrichment of CGC-TSGs/OGs and DORGE-predicted novel TSGs/OGs in hub genes in BioGRID network. (**D**), Enrichment of various gene sets or epigenetic and mutational patterns in hub genes in BioGRID network. (**E**), Enrichment of CGC-TSGs/OGs and DORGE-predicted novel TSGs/OGs in hub genes in the PharmacoDB gene-drug network. (**F**), Enrichment of various gene sets or epigenetic and mutational features in hub genes in the PharmacoDB gene-drug network. Hub genes are defined as the genes with the top 5% highest degree in the BioGRID or PharmacoDB network. To generate comparable *P*-values, the gene number in different gene categories was normalized to 200. HKG: Housekeeping gene; Broad H3K4me3: Genes with H3K4me3 length >4,000; ER: Epigenetic Regulator. *P*-values for the differences between indicated gene categories were calculated by the right-sided Wilcoxon rank-sum test.

It was previously shown that somatic alterations often occur at PPI network hub genes in cancer (*53*), and these hub genes are typically essential genes. We, therefore, investigated the enrichment of cancer driver genes in the hub genes—the 978 genes (top 5%) with the highest degrees in the BioGRID PPI network. We found that all TSGs, OGs, and dual-functional genes (including CGC genes and DORGE-predicted novel genes) are enriched in the hub genes (Fig. 4C). Interestingly, the CGC and novel dual-functional genes are the most enriched (Fig. 4C). We also analyzed the enrichment of ten functional gene sets. Among these gene sets, we found that the genes with high missense o/e constraints (highest top 5%), the essential genes from the OGEE database, and the ER genes are most enriched in the hub genes (Fig. 4D). Previous literature has not reported any connection between ERs and PPI hub genes, and our finding strengthens the critical roles of ERs. We also found that the genes with broad H3K4me3 peaks are significantly enriched, to a similar degree as the housekeeping genes (HKGs), in the hub genes (Fig. 4D).

### Epigenetic regulator genes act as backbones in gene-drug networks

Cancer driver gene prediction is the basis for the development of anti-cancer drugs and personalized cancer treatments. We, therefore, explored possible gene-drug relationships of DORGE-predicted cancer driver genes using the PharmacoDB, a gene-drug network constructed from comprehensive high-throughput cancer pharmacogenomic datasets. In the subnetwork containing CGC genes and DORGE-predicted novel genes, we found that these cancer driver genes are densely connected to anti-cancer drugs (Fig. S5D). Similar to our observation from the PPI network, we found that TSGs and OGs, including CGC genes and DORGE-predicted novel genes, exhibit significantly denser connections to drugs than NGs do (Fig. S5E).

We then identified the top-ten drugs with the largest numbers of connected genes in the PharmacoDB gene-drug network. Among these ten drugs, the top one is doxorubicin, a well-known chemotherapeutic agent, and the other nine drugs are also known anti-cancer drugs (Fig. S5F). We next identified 979 genes (top 5%) with the highest degrees in the gene-drug network as hub genes and found that DORGE-predicted novel driver genes are enriched in these hub genes (Fig. 4E). We also analyzed the enrichment of ten functional gene sets in these hub genes. Unlike their enrichment in our previously defined PPI network hub genes (Fig. 4D), the essential genes and the HKGs are not enriched in these gene-drug network hub genes (Fig. 4F), an expected result as their expression is required for normal cells and they are unlikely to be viable drug targets for cancer treatment. In contrast, we still observed the enrichment of three functional gene sets—the genes with high missense o/e constraints (highest top 5%), the ER genes, and the genes with broad H3K4me3 peaks—in the gene-drug network hub genes (Fig. 4F). Together with our PPI analysis, we conclude that the genes in these three functional gene sets may be potential actionable drug targets. To the best of our knowledge, there has been no report on the enrichment of the ER genes in gene-drug network hub genes. Our results from PPI and gene-drug network analysis emphasize the importance of studying the ER genes as potential drug targets.

### Identification of candidate anti-cancer drugs from public transcriptomic data

A bottleneck in novel anti-cancer drug discovery is an efficient selection of potential molecular targets for a drug/compound or its derivatives. Ideal anti-cancer drugs are those that upregulate TSGs and/or downregulate OGs. We used the CRowd Extracted Expression of Differential Signatures (CREEDS) data (*54*) to explore the relationship between CGC and DORGE-predicted genes and anti-cancer drugs (Data file S1). We identified 68 proven or potential anti-cancer drugs/compounds that were associated with 68 target genes meeting the filtering criteria (limma *Q*-value < 0.05 and fold-change > 2) from the CREEDS data (Fig. S6). Notably, 54 (79.41%) of the 68 genes are DORGE-predicted novel TSG or OG genes.

Recent pharmacological efforts suggested that drugs/compounds actionable toward more than one gene or molecular pathway are preferable for repurposing (*55*), and it is common for existing drugs to be later repurposed as anti-cancer drugs. For example, Dexamethasone was previously classified as a corticosteroid but later repurposed for cancer treatment. Among the 68 drugs/compounds we identified, 30 are anti-cancer and chemotherapy drugs (Fig. S6, bottom), 23 have only been tested in laboratories and are not yet in clinical trials, and 15 have not been tested in cell lines (Fig. S6, bottom). Of the 38 drugs/compounds not yet confirmed in anti-cancer clinical trials, many have been proven to treat other diseases. Overall, our results indicate that they are potential drugs for cancer treatment.

## Discussion

In this paper, we developed a machine-learning tool DORGE for identifying cancer driver genes by integrating genetic and epigenetic features. Our development is the first effort that goes beyond the use of tumor genetic alterations for cancer driver prediction, and it was motivated by our previous studies that found specific epigenetic patterns associated with TSGs or OGs (*19, 21*). Although experimental validation is needed for further studies, our computational evaluation verifies that the novel cancer driver genes predicted by DORGE resemble known cancer drivers in multiple aspects and have promises to be potential therapeutic targets. In particular, the top-ranked novel cancer driver genes, especially those regulated by epigenetic mechanisms, warrant further detailed investigation.

Cancer driver genes that are infrequently mutated in cancer are often indistinguishable from passenger genes with random mutations in genome sequencing data. Such random mutations may result from technical reasons including tumor DNA contamination, sequencing depth, and mutation calling failure (*56*). Therefore, infrequently-mutated cancer driver genes are hardly detectable by the methods based on the mutational background model (MutSigCV (*57*)) or the functional impact model (OncodriveFML (*58*), OncodriveFM (*34*)) and OncodriveCLUST (*59*)). However, these genes may be identified through the integration of epigenetic, phenotypic and genomic data.

In previous studies, various non-mutational datasets have been used in cancer driver gene identification; however, unlike DORGE, existing work only used few or several non-mutational features extracted from these datasets (*7, 50, 51, 57, 60, 61*). For example, MutSigCV used DNA replication timing and cell line expression data (*57*); ActiveDriver used phosphorylation site information (*61*); 20/20+ used multi-species conservation, mutation pathogenicity scores, and replication timing (*7*). PPI networks and pathway knowledge have also been used to identify cancer driver genes (*50, 51*); however, these studies were biased toward well-studied genes/pathways and thus may overlook quite many genuine cancer driver genes. In contrast to all these studies, DORGE leverages epigenetic information without any bias towards gene selection to predict cancer driver genes, and this innovation makes DORGE outpower these existing work in discovering novel cancer driver genes.

We further note that the capacity of DORGE in predicting TSGs and OGs separately allows DORGE to identify novel dual-functional cancer driver genes. This is advantageous given that more and more dual-functional cancer driver genes have been identified in the literature. In this study, we found a unique property of dual-functional cancer driver genes: they have more connecting partners in PPI and drug-gene networks than other driver genes have. This property, to our knowledge, was not previously reported. In fact, several novel dual-functional genes predicted by DORGE drew our attention. For example, *PTEN* (*Phosphatase and tensin Homolog*), a protein phosphatase, is commonly regarded as a TSG; however, DORGE predicted it as an OG as well. We found that, indeed, oncogenic roles were reported for *PTEN* in a few studies. One explanation for the dual-functional roles of *PTEN* is that its oncogenic effect depends on the positions of mutations (*62*). We confirmed this by analyzing the mutation patterns of *PTEN* and found one pattern as the classic OG mutation pattern with most substitutions in p.R130 (*63*). In DORGE, further updates can quantify the dual-functional roles (i.e. the relative chance of being TSGs or OGs) of dual-functional genes.

While we have already found dozens of non-mutational features that contribute significantly to the predictive power of DORGE, many CGC genes remain undetected by DORGE (Fig. 2G and H). A possible reason is the missingness of other factors or mechanisms that regulate cancer driver genes. Fortunately, the continual increase in functional genetic and epigenetic data will provide a lasting opportunity to improve and fine-tune cancer driver gene prediction methods. In future studies, we can perform lineage-specific rather than pan-cancer prediction and extend DORGE to predicting long non-coding genes, as many features used in DORGE are not restricted to protein-coding genes. In addition, further work is needed for a better understanding of the reasons underlying the phenomena such as ancient phyletic ages of cancer driver genes and enrichment of cancer driver genes at PPI and gene-drug network hubs.

In summary, this study highlights the integration of epigenetic data to achieve a more comprehensive prediction of cancer driver genes. DORGE will serve as an essential resource for cancer biology, particularly in the development of targeted therapeutics and personalized medicine for cancer treatment.

## Materials and Methods

### Experimental Design

In this paper, we propose DORGE, a machine-learning framework incorporating a large number of features to discover TSGs/OGs (Fig. S2). First, we used CGC v.87 genes and NGs as the training genes to predict TSGs and OGs separately from 75 candidate features by logistic regression with the elastic net penalty, and the resulting two classifiers are DORGE-TSG and DORGE-OG. Next, we used five-fold cross-validation to evaluate DORGE. We also analyzed the benefit of introducing epigenetic features based on KEGG enrichment and evaluated DORGE based on several genomic and functional genomic datasets. Lastly, we showed the enrichment of dual-functional genes predicted by DORGE in hub genes in PPI and gene-compound networks.

### Gene annotation

All gene annotations, genomic and functional genomic datasets were downloaded from hg19 genome version or processed to hg19 if from other genome versions. Genome version conversion was done using the LiftOver program (https://genome.ucsc.edu/cgi-bin/hgLiftOver). HUGO Gene Nomenclature Committee (HGNC) annotation (https://www.genenames.org/) was used for gene annotation. The gene annotation can be found in the Data file S1. Promoters were defined as the regions from the upstream 1,000 bp to downstream 500 bp of Transcription Start Sites (TSSs), while gene-body regions were defined as the regions from downstream 500 bp of TSSs to Transcription Termination Sites (TTSs).

### Datasets used in this study

#### Somatic mutation datasets

The somatic mutation dataset used in this study was derived from the TCGA (*6*) website (https://portal.gdc.cancer.gov/) and the Catalogue Of Somatic Mutations in Cancer (COSMIC), v86 (*5*). These two datasets were combined to help increase the mutational information of infrequently mutated genes. Duplicate tumor samples present in more than one dataset were excluded. The final dataset used for the calculation of mutation-related features contained 5,700,484 mutations from more than 30 tumor types. Hypermutated tumor samples with >2,000 mutations were excluded from this dataset. The population genetic dataset for evaluating features, such as loss-of-function (LoF) intolerance, was downloaded from The Genome Aggregation Database (gnomAD) (https://storage.googleapis.com/gnomad-public/release/2.1.1/constraint/gnomad.v2.1.1.lof_metrics.by_gene.txt.bgz) (*64*). Additional details regarding features calculation can be found in the Data file S1.

#### Epigenetic datasets

We downloaded all peak BED files (hg19) for tri-methylation on histone H3 lysine 4 (H3K4me3) and other representative histone modifications from the ENCODE project (https://www.encodeproject.org/). The full file names and download links are listed in the Data file S1. The gene-body canyon annotation file (*65*), including DNA methylation information, was obtained from a previous study (*21*), which were based on TCGA whole-genome bisulfite sequencing (WGBS) data. The data for calculating promoter and gene-body cancer-normal methylation difference was also downloaded from the level 3 methylation data from the COSMIC website (v.90). Repli-seq BAM datasets were downloaded from the ENCODE project website, and the featureCounts program (http://subread.sourceforge.net/) was used to assign BAM reads to gene-bodies. Read counts were normalized based on the sequencing depth of the BAM files, and the normalized read numbers were used to calculate the replication timing S50 score (*66*). This score, which determines the median replication timing, was calculated by a tool available from a previous study (*66*). The super enhancer annotation was downloaded from the dbSUPER database (*23*).

#### Other datasets

The level 3 TCGA data, which includes the processed somatic copy number alteration (CNA) and gene expression data, were downloaded from the COSMIC website (v.90) and used without processing. The processed cell proliferation (dependency) scores from 436 CRISPR-treated cell line samples were obtained from the DepMap website (Avana-17Q2-Public_v2) (*24*). For each gene, gene expression was aggregated across samples to obtain the median *Z* score. The phastCons scores were downloaded from the UCSC (http://hgdownload.cse.ucsc.edu/goldenPath/hg19/phastCons46way/). The dataset including the gene damage index (GDI), Primate *dN*/*dS* score, Residual Variation Intolerance Scores (RVIS) percentile, non-coding Residual Variation Intolerance Scores (ncRVIS), non-coding Genomic Evolutionary Rate Profiling (ncGERP), family member count and gene age features were downloaded from https://github.com/RausellLab/NCBoost (*22*). The dataset is gene-centric, and no further processing was done.

### Curation of TSG, OG, and NG training sets

The training set contained 242 high-confidence TSGs and 240 high-confidence OGs without overlapping from the v.87 CGC database on the COSMIC website, as well as 4,058 NGs obtained as follows. The initial set of NGs was obtained from Davoli *et al.* (*9*). However, this initial set is likely to include mislabeled genes. To address this, those that overlap with the following gene lists (June 18, 2020) were excluded from this initial NG set: (I) Candidate Cancer Gene Database (*67*), (II) CancerMine (*32*), (III) a cancer gene list compiled by Chiu et al. (*68*), (IV) the genes (OncoScore > 21.09) in OncoScore database (*69*), and (V) allOnco Cancer Gene List (v3 Feb 2017; http://www.bushmanlab.org/links/genelists). The final training gene sets are available at Data file S2.

### Candidate mutational features

The candidate mutational features were previously defined by Davoli *et al.*(*9*) and Tokheim *et al.* (*7*). In addition to these features, other gene-centric features were also collected. Features were categorized into the following classes: ‘Genomics’, ‘Mutation’, ‘Epigenetics’ and ‘Phenotype’, and additional details regarding these features can be found in Data file S1. The feature IDs mentioned below correspond to Data file S1.

Features 1–20 were quantified based on the combined mutation data using the script provided by Davoli *et al.* (*9*). Further information for these features can be found in their paper (*9*). For features 1, 5, and 6 in Data file S1 that quantify the density of different categories of mutations within genes, only the coding sequence (CDS) length (per kb) of each gene is considered. For mutational features 8–15 and 28 that include ratios, a pseudo count estimated as the median of each feature across all genes was added, as described by Davoli *et al.* (*9*).

Features 11–15 rely on the functional effects of missense mutations, including high functional impact (HiFI) or low functional impact (LoFI) (*9*). The PolyPhen-2 Hum-Var prediction model was used to estimate the functional effects of missense mutations and to classify them as either high functional impact (HiFI) or low functional impact (LoFI) (*9*), based on the probability of functional damage as estimated by the PolyPhen-2 HumVar algorithm. Features based on HiFI and LoFI include: 1) benign mutations: silent and LoFI missense mutations; 2) LoF mutations: nonsense and frameshift mutations; and 3) HiFI missense mutations (damaging missense mutations). PolyPhen-2 scores (Feature 16) were calculated by the PolyPhen-2 web server (http://genetics.bwh.harvard.edu/pph2/)(*70*). The missense MGAentropy scores (Feature 33), which also measure the multi-species conservation of missense mutation sites, were also calculated by the CRAVAT tool (*71*).

Other mutation types include splicing/total mutations (Feature 19) and inactivating fraction (Feature 27). Splicing mutations are those that affect splicing sites; >95% of splicing mutations are in the first two positions at donor or acceptor sites. Inactivating mutations include indel frameshift, splice site, translation start site, and nonstop mutations. Features 21-29 that were introduced in Tokheim et al.’s paper were quantified based on our revised version of the script provided by Davoli *et al.* (*9*), given that these features can be quantified in a similar way to that for Features 1–20. The lost start and stop fraction (Feature 26) was defined as the fraction of the translation start site, and nonstop mutations in total mutations. The recurrent missense fraction (Feature 23) was defined by missense mutations occurring more than one patient sample.

Features 42–46 are population genetics-based mutational features. For LoF constraints, three categories of tolerance to LoF mutations were defined by gnomAD: null (LoF mutations are fully tolerant), recessive (heterozygous LoF mutations are tolerant), and haploinsufficient (heterozygous LoF mutations are intolerant). The probability of the three types of mutations can also be obtained from the dataset (Features 42–43), or be derived based on simple calculation (Sum of the probability of three categories of intolerance equals one). A probability of being LoF intolerant (pLI) score was initially introduced to determine the likelihood that a given gene is intolerant of LoF mutations. The difference between LoF o/e and pLI is explained at https://blog.limbus-medtec.com/how-to-use-gnomad-v2-1-for-variant-filtering-d7d2a7ee710a. For synonymous, missense, and LoF mutations (Features 44–46), a signed *Z* score to describe the deviation of observation from expectation (o/e) was obtained from the gnomAD dataset. Higher *Z* scores indicate intolerance to variation or increased constraint, whereas lower *Z* scores indicate tolerance to variants.

### Candidate epigenetic features

In addition to genetic data, epigenetic data have been shown to be associated with cancer driver genes. Here, we used the peak length and height to characterize histone modifications. We also used cancer– normal methylation difference to characterize gene promoter and gene-body methylation in cancer and normal samples. These potentially useful features (Features 39–40 and 54–75) were previously used in epigenetics studies, but to what extent these features are useful in predicting cancer driver genes are not systematically evaluated. The histone modification BED files were processed based on our previously published procedures (*19*). Briefly, adjacent peaks were merged when peaks are within 3-kb by the merge command from bedtools (https://bedtools.readthedocs.io/). Peaks overlapping with the longest transcript of a gene at least 50% of peak length were assigned to that gene by bedmap function in the BEDOPS tool (https://bedops.readthedocs.io/) with the following parameters: --max-element --echo --fraction-map 0.5 --delim ‘\t’ --skip-unmapped. Features of “Mean peak length” were calculated based on the merged peaks. For features of “height of peaks”, the maximum signal values (7th column in BED 6+4 narrow peak files used in ENCODE) were used. Promoter and gene-body cancer–normal methylation difference features (Features 39 and 40) were defined by the mean methylation level in cancer samples (Beta Value column in the dataset) minus that in normal samples (Avg Beta Value Normal column in the dataset) based on COSMIC 450K methylation data. 450K probes were mapped to genes according to genomic coordinates (hg19). The gene-body canyon cancer/normal methylation ratio feature (Feature 41) was inspired from a previous study (*21*). The ratio value was determined by the mean methylation level in cancer samples devided by that in normal samples in TCGA WGBS methylation data. To make “Gene-body cancer– normal methylation difference” (Feature 40) and “Gene-body canyon cancer/normal methylation ratio” (Feature 41) available to all genes, genes without applicable feature values were imputed as 0. Genes were linked to gene-body canyons by BEDOPS with the same parameters as shown above. The difference between Feature 40 and 41 is that Feature 41 is only available to genes with gene-body methylation canyons defined by a previous study using TCGA WGBS data (*21*), while Feature 40 is available for all genes with 450K probes. We previously used TCGA WGBS data to define Feature 41 because WGBS methylation data has a significantly higher resolution than 450K methylation data, while we were unable to identify large DNA methylation canyons using COSMIC 450K data. For feature 34, Repli-seq BAM datasets were quantified by the featureCounts program (http://subread.sourceforge.net/) to assign BAM reads to gene-bodies. Read counts were normalized based on the sequencing depth of the BAM files, and the normalized read numbers were used to calculate the S50 score (*66*). Early replication timing (Feature 34) was quantified by the S50 score. All bam data are assigned to different cell cycle stages (G1, S1, S2, S3 and S4) for the S50 score calculation. This score, which determines the median replication timing (from 0–1), was calculated based on the algorithm proposed by a previous study (*66*). A S50 score that closes to 0 means early replication timing, whereas a S50 score that closes to 1 means late replication timing. Super enhancer percentage (Feature 38) was calculated as the percentage of cell lines in which super enhancers are associated with any transcripts of genes.

### Other candidate features

Feature 29 (log gene length) was defined as the log_2_ transformed length of the maximum transcript of a specific gene based on the ENSEMBL GTF annotation file. Feature 30 (log CDS length) was obtained from the COSMIC mutation files and supplemented by the ENSEMBL GTF annotation file, and then log_2_ transformed. Features 31 is CNA deletion percentage which was calculated based on column 17 (Mut Type: gain or loss) in the original dataset (CNA amplification percentage can be calculated by 1 - CNA deletion percentage). The Variant Effect Scoring Tool (VEST) scores (Feature 35), which indicate missense pathogenicity for each mutation, were calculated by CRAVAT. Gene expression Z score (Feature 36) was used to quantify the gene expression based on the “Regulation” column in the original data. The exon conservation (phastCons) score (Feature 32) that is based on the average phastCons score for maximum transcripts of genes was also calculated by CRAVAT. Feature 37 (Increase of cell proliferation by CRISPR Knock-down) was calculated based on the cell proliferation scores in the CRISPR screening data. A lower cell proliferation for a gene in a cell line means that the gene is more likely to essential to the cell line. A score of 0 means nonessential, whereas a score of −1 means essential.

Features 47–53 are evolution-based features, including GDI (Mutational damage that has accumulated in the general population), Primate *dN*/*dS* score (Ratio between the number of nonsynonymous substitutions and the number of synonymous substitutions), RVIS percentile (High RVIS percentiles reflect genes highly tolerant to variation), ncRVIS, ncGERP, family member count (Number of human paralogs for each gene), and the gene age (Time of evolutionary origin based on the presence/absence of orthologs in vertebrates). Genes with higher GERP scores are more constrained. ncRVIS is a measure of deviation from the genome-wide variants found in non-coding sequences of genes (*46*). A negative ncRVIS score indicates less common non-coding variation than predicted. In ncRVIS and ncGERP, the non-coding regions were defined as the untranslated regions (UTRs) as well as non-exonic 250 bp upstream of TSSs.

### Training of DORGE-TSG and DORGE-OG

The elastic net is a penalized regression method that can select a limited number of features that contribute to the response. Similar to the lasso, the elastic net selects features by shrinking some of the coefficients to be zero; the remaining features with nonzero coefficients are considered to have larger effects on the response and thus are selected and kept in the model. The main advantage of the elastic net over the lasso is that in case of collinearity the elastic net simultaneously selects a group of colinear features whereas the lasso tends to select only one feature from the group. (The simultaneous selection of collinear features is desired because, in the extreme situation where these collinear features are exactly identical, the regression method should assign equal coefficients to these features.) Therefore, we chose the elastic net over the lasso because we observed high collinearity among the original list of 75 features.

Specific to DORGE, we used logistic regression with the elastic net penalty to train two binary classifiers for predicting TSGs and OGs, and these classifiers were referred to as DORGE-TSG and DORGE-OG. We used the R function glmnet from the R package glmnet (https://cran.r-project.org/web/packages/glmnet/index.html). The *λ* tuning parameter was selected by 5-fold cross-validation using the function cv.glmnet from the same R package, while the *α* parameter, which balances the lasso and ridge penalties, was set to the default value 0.5.

For every gene, DORGE-TSG predicted it with a probability of being a TSG, and this probability is defined as the gene’s TSG-score. The OG-scores are defined similarly by DORGE-OG for all genes. Having two separate binary classifiers, one for detecting TSG and the other for detecting OG, DORGE is able to detect dual-functional genes.

The codes for training DORGE-TSG and DORGE-OG and obtaining predicted TSGs and OGs is available at https://github.com/biocq/DORGE. An online video that explains the code is available at https://www.youtube.com/watch?v=Pk8ZqoHK8zk.

### Precision-Recall Curve analyses

Precision-recall curve (PRC) analyses were performed using the R PRROC. The AUPRCs were calculated using TSG-scores and OG-scores by the pr.curve function in the package.

### Thresholds on TSG-scores and OG-scores

We used in-house code available in our DORGE GitHub repository to find the cutoffs on TSG-scores and OG-scores such that the population false positive rates (type I errors; for TSG prediction, the false positive rate is the conditional probability of misclassifying an NG as a TSG) were controlled under 1%. The code was an implementation of the Neyman-Pearson classification umbrella algorithm (*25*).

### Gene sets, genomic and functional genomic datasets used for characterization and evaluation of DORGE-predicted novel TSGs and OGs

The gene lists and datasets that we used to evaluate our DORGE-predicted novel TSGs/OGs are as follows: (I) CGC gold-standard gene list. The CGC is a widely used gold-standard list of cancer-related genes. We used the CGC v.87 gene list as the testing gene set while excluding those in v.77 CGC gene list to evaluate the performance of our prediction. (II) ATAC-Seq data. ATAC-Seq data were taken from pan-cancer peak calls from Data S2 in Corces et al.’s paper (*39*). (III) Epigenetic regulators (ERs). The ER gene list comes from a recent study focused on the characterization of ERs (*40*) and the EpiFactors database (*72*), after removing the genes that function only as TFs. (IV) Candidate TSGs identified by Sleeping Beauty insertional mutagenesis. The inactivating pattern gene list was downloaded from the Sleeping Beauty Cancer Driver Database (SBCDDB) (*42*). This database contains cancer driver genes that were identified by Sleeping Beauty insertional mutagenesis in tumor models. For the evaluation of DORGE-predicted novel TSGs, only genes with an inactivating pattern in the SBCDDB were kept, resulting in 1,211 genes. (V) Survival data. Survival data were downloaded from OncoLnc website (*44*). (VI) shRNA screening data. The gene-centric shRNA screening data (v2.4.5) were taken from the Achilles project (*43*). (VII) Evolutionary conservation data. For evolutionary conservation, we used phyloP scores that measure non-neutral substitution rates based on multi-species alignments. The phyloP data were downloaded from the University of California, Santa Cruz (UCSC); http://hgdownload.cse.ucsc.edu/goldenPath/hg19/phyloP46way). We computed the average −log(phyloP) and phastCons score for each gene by averaging the base-pair-level conservation values for every position in each gene. (VIII) Phyletic age. We downloaded the precomputed phyletic age gene lists in human and measured enrichment of our predicted genes within the gene sets from different phyletic ages (i.e., Eukaryota, Metazoa, Chordata, and Mammalia) from the Online GEne Essentiality (OGEE) database (*48*). (IX) The BioGRID v3.5.183 data were downloaded from the website, https://thebiogrid.org/. Biological network related metrics can be calculated by the Cytoscape software (*73*). Additional information on the network metrics can be found in the Supplementary Text. (X) The PharmacoDB (*74*) gene-drug network data were downloaded from https://pharmacodb.pmgenomics.ca/. (XI) Housekeeping genes (HKGs). We downloaded an HKG gene list from https://www.tau.ac.il/~elieis/HKG/, which includes 3,804 HKGs. (XII) Essential genes. The essential and nonessential gene lists were also downloaded from the OGEE database. To shorten this list, we limited our essential gene set to those with >2 in entries of the OGEE database, resulting in 2,340 definitive essential genes. Non-essential genes that overlap with essential genes were removed, resulting in 11,990 non-essential genes. (XIII) The drug response data were downloaded from Drug Gene Budger (*54*). Only significant drug-gene relationships (*Q*-value < 0.05 and fold-change > 2) were selected from the CRowd Extracted Expression of Differential Signatures (CREEDS) data collections downloaded from the Drug Gene Budger (DGB) database (*54*).

### Gene-set enrichment analysis

Kyoto Encyclopedia of Genes and Genomes (KEGG) enrichment analyses were performed using Enrichr (*75*) for DORGE and DORGE variant predicted novel genes.

### Protein-protein interaction network module analysis

For DORGE-predicted novel genes and CGC genes, PPI module analysis was performed by Metascape (*76*). Networks contain proteins that display physical interactions with at least one other protein in the list. For networks containing 3 to 500 proteins, the Molecular Complex Detection (MCODE) algorithm (*77*) was applied to identify densely connected network modules.

### Statistical Analysis

One-sided Wilcoxon rank-sum test was used when comparing different categories of genes. Gene enrichment analyses were performed in R, using one-sided Fisher’s exact test (fisher.test function in R). *P*-values of Spearman correlation were calculated by Test for Association/Correlation Between Paired Samples (cor.test function in R). Binomial test was used to test the enrichment of dual-functional genes in network hub genes.

## Acknowledgments

We would like to thank potential reviewers for their insightful suggestions and comments on this paper.

## Author contributions

Conceptualization: W.L., J.L. and J.J.L.; methodology: J.J.L. and J.L.; software: J.J.L. and J.L.; writing: J.J.L., J.L., Y.C., X.G. and W.L.; data analysis: J.L., J.J.L., J.S., F.P. and X.G.; supervision: W.L. and J.J.L.

## Data and materials availability

The open source codes for DORGE are freely available at https://github.com/biocq/DORGE or upon request, along with the data and scripts to reproduce the results of the paper.

