## Supplementary file for "DORGE: Discovery of Oncogenes and Tumor SuppressoR Genes Using Genetic and Epigenetic Features"

**Affiliations**

The PDF file includes:

Supplementary Text

Fig. S1. Features that discriminate tumor suppressor genes (TSGs) from oncogenes (OGs).

Fig. S2. Flowchart for the illustration of DORGE method.

Fig. S3. Performance of DORGE is better than the TUSON and 20/20+ approaches for evaluation on CGC genes.

Fig. S4. Evaluation of DORGE-predicted novel TSG/OGs by independent functional genomic and genomic datasets.

Fig. S5. Dual-function TSG/OGs are more highly enriched in the BioGRID protein-protein interaction (PPI) and the PharmacoDB gene-compound networks.

Fig. S6. Heatmap of fold changes in expression for DORGE-predicted novel and CGC genes that are up- or down-regulated by drug perturbations.

Legends for data files S1 and S2

Other Supplementary Material for this manuscript includes the following:

Data file S1 (Microsoft Excel format). Excel spreadsheet detailing the supplemental data including the candidate features, Spearman correlation table for features, histone modification data information, gene annotation, the DORGE predicted TSGs/OGs without occurring in cancer driver databases as well as the associated survival hazard ratios (HRs), and also the gene expression responses data for different drugs/compounds.

Data file S2 (Microsoft Excel format). Excel spreadsheet detailing AUPRC values of eight classification algorithms under three class ratios, the feature groups, the DORGE prediction results and training gene annotations, evaluation of DORGE driver gene prediction on the OncoKB gene set, evaluation of DORGE TSG/OG prediction on the CGC gene list, the features used in different cancer driver prediction studies as well as the training feature profile.

**Supplementary Text**

**Network degree, betweenness, and closeness centrality**

In the BioGRID network, protein-protein interactions (PPIs) are illustrated with undirected graphs whose nodes represent proteins, and edges between two nodes represent existing interactions between the connected proteins. We downloaded the human PPI network from BioGRID v.3.5.183. Interactions were reduced to include only nonredundant interacting proteins in our network visualization. Betweenness centrality, which measures the centrality of each gene in the network, was calculated as follows. Given a network with *n* genes,
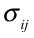
is defined as the total number of shortest paths between gene *i* and gene *j*, and
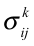
 is defined as the total number of
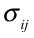
that passes through gene *k*. The betweenness centrality for gene *k* is defined as:
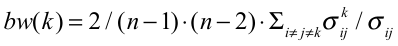
. Closeness centrality, which is also often used to describe the centrality of a biological network, was calculated as the reciprocal of the sum of the length of the shortest paths between the gene and all other genes in the network. The closeness centrality is defined as: *C*(*x*) *= N /* Σ*_y_ d*(*y*, *x*), where *N* is the number genes in the network, and *d*(*y*, *x*) is the distance between genes *y* and *x*. Thus, the more central a gene, the closer it is to all other genes. The network degree metrics for PharmacoDB gene-drug/compound network was also calculated in the same way.

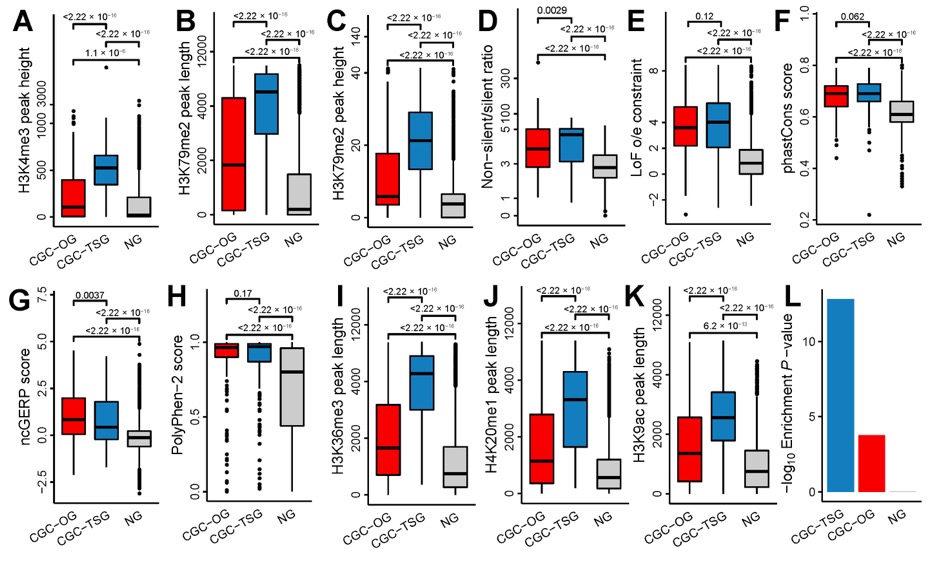

**Fig. S1.** **Features that discriminate tumor suppressor genes (TSGs) from oncogenes (OGs).** Box plots showing the distribution of (**A**), Tri-methylation of histone H3 lysine 4 (H3K4me3) peak height, (**B**), Di-methylation of histone H3 lysine 79 (H3K79me2) peak length, (**C**)**,** Di-methylation of histone H3 lysine 79 (H3K79me2) peak height, (**D**), Non-silent/silent ratio, (**E**), LoF o/e constraint, (**F**), phastCons score, (**G**), non-coding Genomic Evolutionary Rate Profiling (ncGERP) score, (**H**), PolyPhen-2 score, (**I**), Tri-methylation of histone H3 lysine 36 (H3K36me3) peak length, (**J**), Mono-methylation of histone H4 lysine 20 (H4K20me1) peak length, (**K**), Histone H3 lysine 9 acetylation (H3K9ac) peak length. Boxplots show the lower, median, and upper quartile of values, and lines extend to 1.5 times of the interquartile range among the Cancer Genomics Consortium (CGC)-OG, CGC-TSG, and neutral gene (NG) sets. (**L**), Enrichment *P*-values based on Fisher’s exact test for genes with broad H3K4me3 peaks (length > 4,000 bp) in the CGC-OG, CGC-TSG, and NG sets.

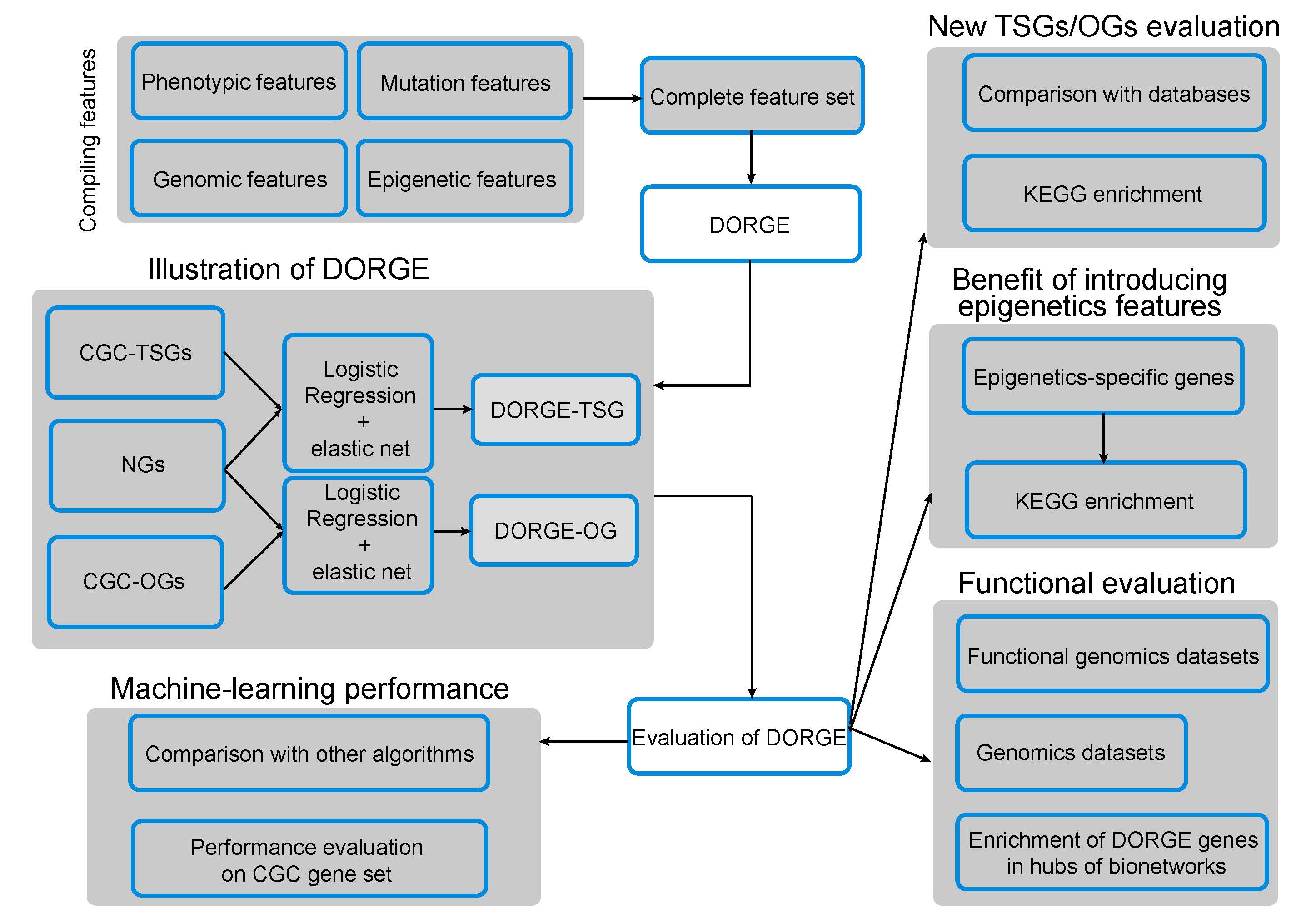

**Fig. S2.** Flowchart for the illustration of DORGE method. The inputs of the method are taken from the public resources including pan-cancer mutation and epigenetic datasets. The training of the DORGE model is based on v.87 CGC TSGs and OGs (excluding dual-function genes) as well as NGs by Logistic regression with elastic net penalty. Predictions of TSGs and OGs are done independently by DORGE-TSG and DORGE-OG model. This method outputs probability as scores for ranking genome-wide genes. Based on the DORGE-TSG and DORGE-OG score, we obtain the DORGE-predicted TSG/OG gene sets based on FPR < 1%. The performance of DORGE for predicting cancer driver genes and TSG/OGs is compared with other algorithms and is evaluated based on CGC and OncoKB gene lists. Top predicted novel TSG/OGs are compared with literature and databases and evaluated by KEGG pathway enrichment. The benefit of epigenetics features is demonstrated by evaluating the contribution of epigenetics features to the prediction and evaluating the integrative model through KEGG enrichment vs. variant model without epigenetics features. The DORGE gene set is then evaluated using several genomics or functional genomics datasets. We also explore the enrichment of DORGE genes in hubs of bionetworks including protein-protein interaction (PPI) and gene/compound-drug networks.

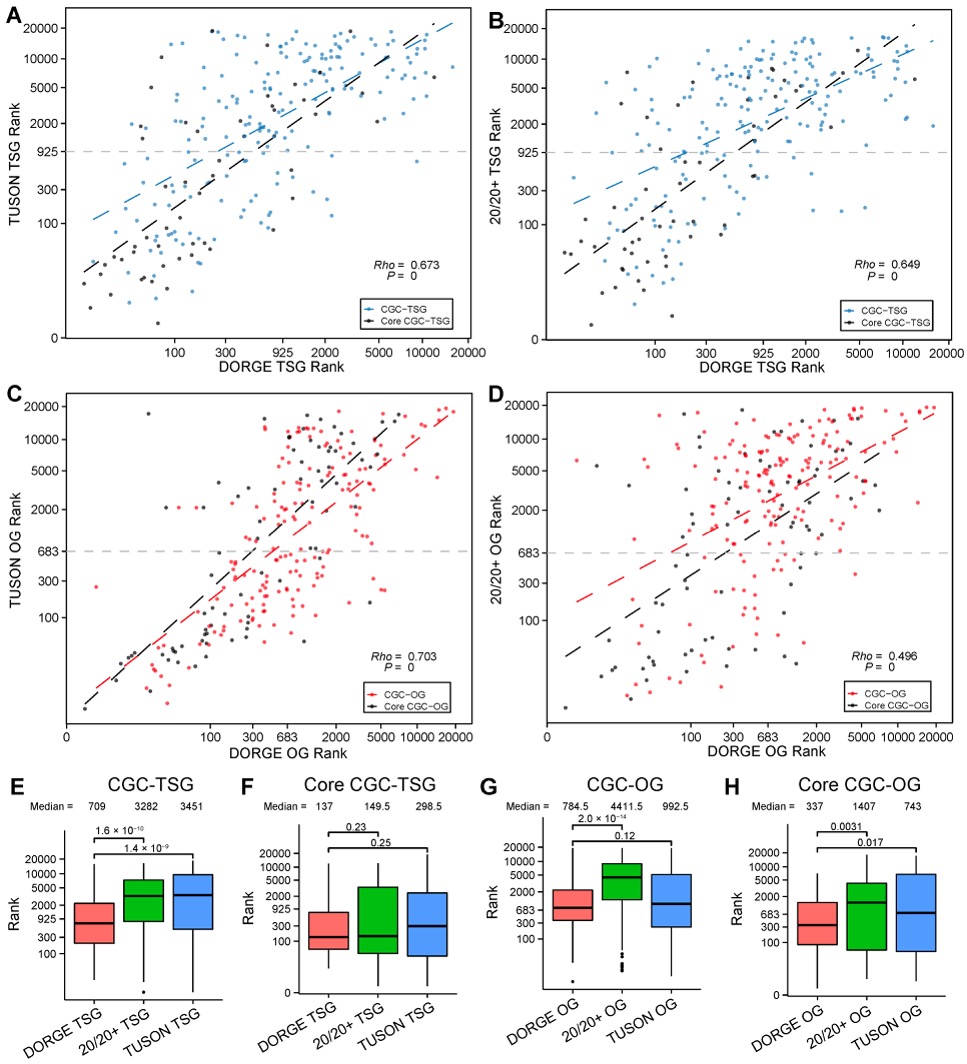

**Fig. S3. Performance of DORGE is better than the TUSON and 20/20+ approaches for evaluation on CGC genes.** (**A**), Scatter plot comparing TSG prediction by DORGE and TUSON using CGC-TSGs. (**B**), Scatter plot comparing TSG prediction by DORGE and 20/20+ using CGC-TSGs. (**C**), Scatter plot comparing OG prediction by DORGE and TUSON using CGC-OGs. (**D**), Scatter plot comparing OG prediction by DORGE and 20/20+ using CGC-OGs. Core CGC genes are defined as the genes from CGC v.77 while excluding CGC-dual-function genes in CGC v.77. Spearman correlation coefficient *Rho* between ranks of genes predicted by different methods in (**A**)–(**D**) was calculated, and *P*-values were calculated by Test for Association/Correlation Between Paired Samples (cor.test function in R). Horizontal gray dashed lines in (**A**)–(**D**) indicate the threshold for DORGE prediction. Linear regression lines by least squares are also indicated for CGC and core CGC genes. (**E**) Boxplots showing the ranks of all CGC-TSG (v.87) predicted by three methods. (**F**) Boxplots showing the ranks of core CGC-TSG (v77) predicted by three methods. (**G**) Boxplots showing the ranks of all CGC-OG (v.87) predicted by three methods. (**H**) Boxplots showing the ranks of core CGC-OG (v.77) predicted by three methods. *P*-values for the differences between indicated gene categories in (**E**)–(**H**) were calculated by the one-tailed Wilcoxon rank-sum test.

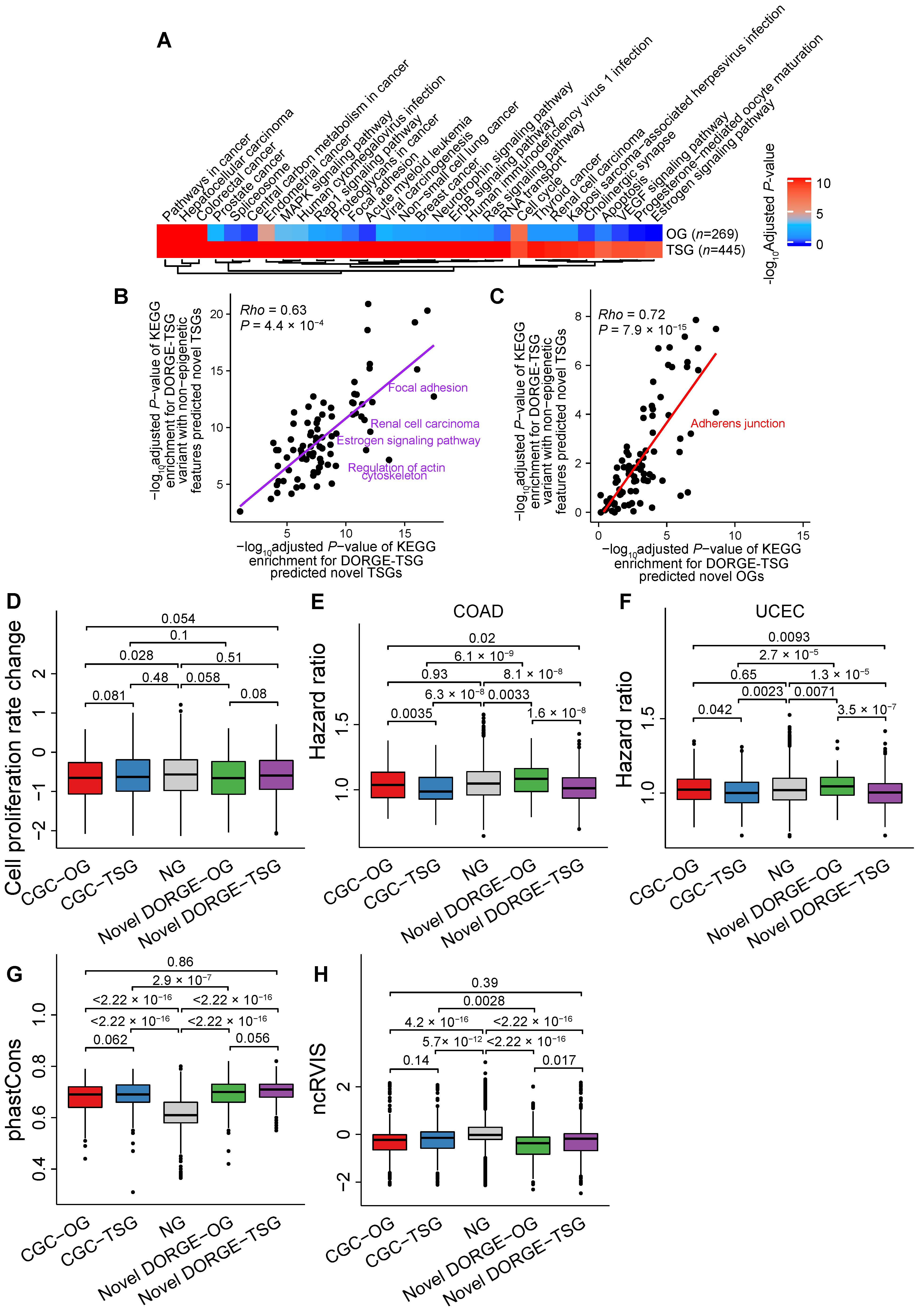

**Fig. S4.** **Evaluation of DORGE-predicted novel TSG/OGs by independent functional genomic and genomic datasets.** (**A**), Kyoto Encyclopedia of Genes and Genomes (KEGG) pathway enrichment analysis for the predicted novel TSGs and OGs that are predicted by DORGE variant with non-epigenetic features. Enrichment analysis was performed by Enrichr (http://amp.pharm.mssm.edu/Enrichr/). Due to space limitation, terms with adjusted *P*-values < 10^-4^ are shown. Besides, terms with adjusted *P*-values 10^8^-fold lower for TSGs than OGs or 10^4^-fold lower for OGs than TSGs are also shown. (**B**), Scatter plot comparing -log_10_ adjusted *P*-values of KEGG enrichment for non-CGC DORGE-TSG predicted novel genes predicted by DORGE-TSG and DORGE-TSG variant with non-epigenetic features. (**C**), Scatter plot comparing -log_10_ adjusted *P*-values of KEGG enrichment for non-CGC DORGE-OG predicted novel genes predicted by DORGE-OG and DORGE-OG variant with non-epigenetic features. Spearman correlation coefficient *Rho* in (**B**)–(**C**) was calculated by the Spearman correlation and *P*-values were calculated by Test for Association/Correlation Between Paired Samples (cor.test in R). (**D**), Boxplot showing the cell proliferation rate change by shRNA-mediated gene knockdown for CGC-TSG/OGs, novel DORGE-TSG/OGs and NGs. A lower score means that a gene is more likely to be dependent in all investigated cell lines. (**E**), Boxplot showing the Cox hazard ratio (HR) score for CGC-TSG/OGs, novel DORGE-TSG/OGs and NGs in Colon adenocarcinoma (COAD) and (**F**), in Uterine Corpus Endometrial Carcinoma (UCEC). (**G**), Boxplot showing the phastCons score for NGs, CGC- TSGs/OGs, and novel DORGE-TSGs/OGs. phastCons conservation scores for the entire exon were calculated from a 46-species phylogenetic alignment from vertebrate, mammals, and primate species and downloaded from the University of California, Santa Cruz (UCSC) (http://hgdownload.cse.ucsc.edu/goldenPath/hg19/phastCons46way/). (**H**), Boxplot showing the non-coding Residual Variation Intolerance Score (ncRVIS) score for CGC-TSGs/OGs, novel DORGE-predicted TSG/OGs, and NGs. ncRVIS is a measure of the deviation from the genome-wide variation found in the non-coding sequence of genes with a similar amount of non-coding mutational burden. A negative ncRVIS score indicates less common non-coding variation than predicted, whereas a positive ncRVIS score indicates more. *P*-values for the differences between indicated gene categories were calculated by the one-tailed Wilcoxon rank-sum test.

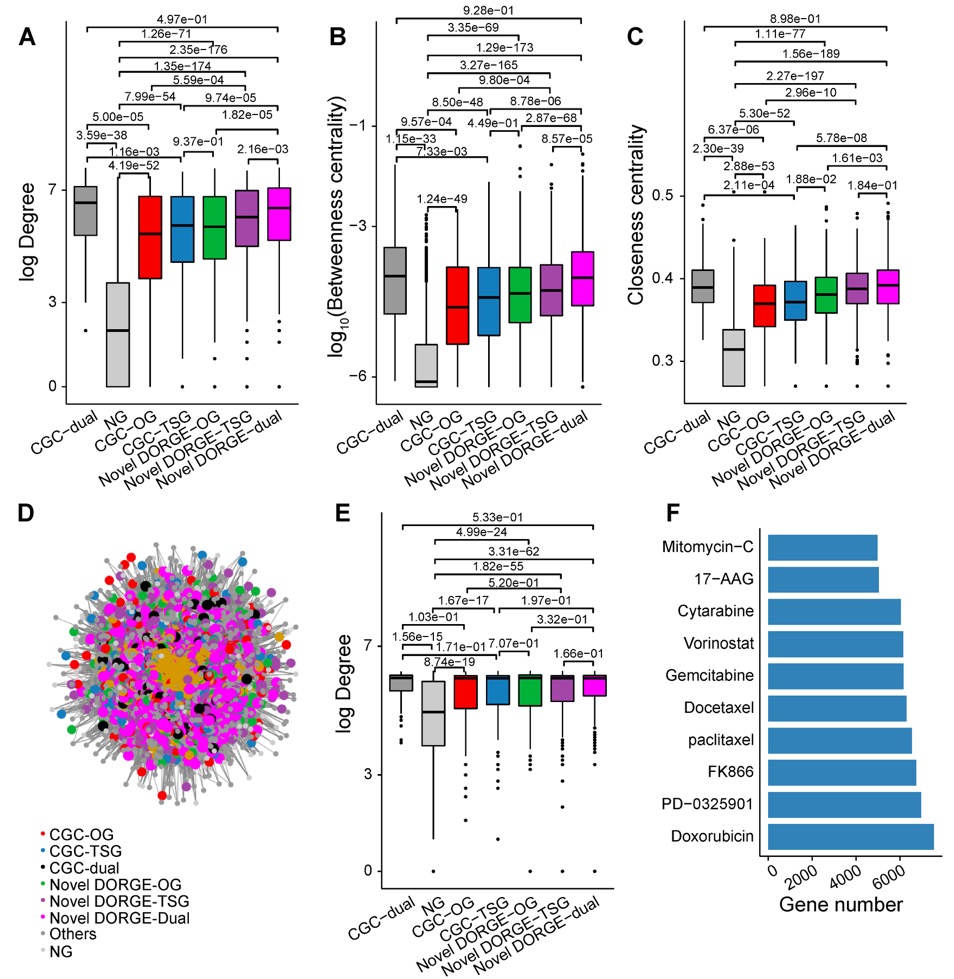

**Fig. S5. Dual-function TSG/OGs are more highly enriched in the BioGRID protein-protein interaction (PPI) and the PharmacoDB gene-compound networks.** (**A**), Plots showing the degree, (**B**), betweenness centrality, and (**C**), closeness centrality of CGC and novel DORGE-TSG/OGs, dual-function genes, and NGs in the BioGRID PPI network. CGC-TSGs/OGs and novel DORGE-TSGs/OGs display similar network metrics in the BioGRID network while CGC- and DORGE-predicted novel-dual-function genes display larger network connectiveness. (**D**), CGC-TSGs/OGs and novel DORGE-TSGs/OGs are densely connected in the gene- drug/compound interaction network. This network was constructed based on the PharmacoDB database, which is a comprehensive *in vitro* drug screen database. Only interactions with an analysis of variance (ANOVA) *P*-values < 0.01 are shown. (**E**), CGC-TSGs/OGs and novel DORGE-TSGs/OGs display similar network degrees in the PharmacoDB network. CGC-dual-function genes and DORGE-predicted novel-dual-function genes are similarly associated with anti-cancer drugs compared to other gene categories. (**F**), Bar plot showing the top ten anti-cancer drugs with the most associated genes in the PharmacoDB database.

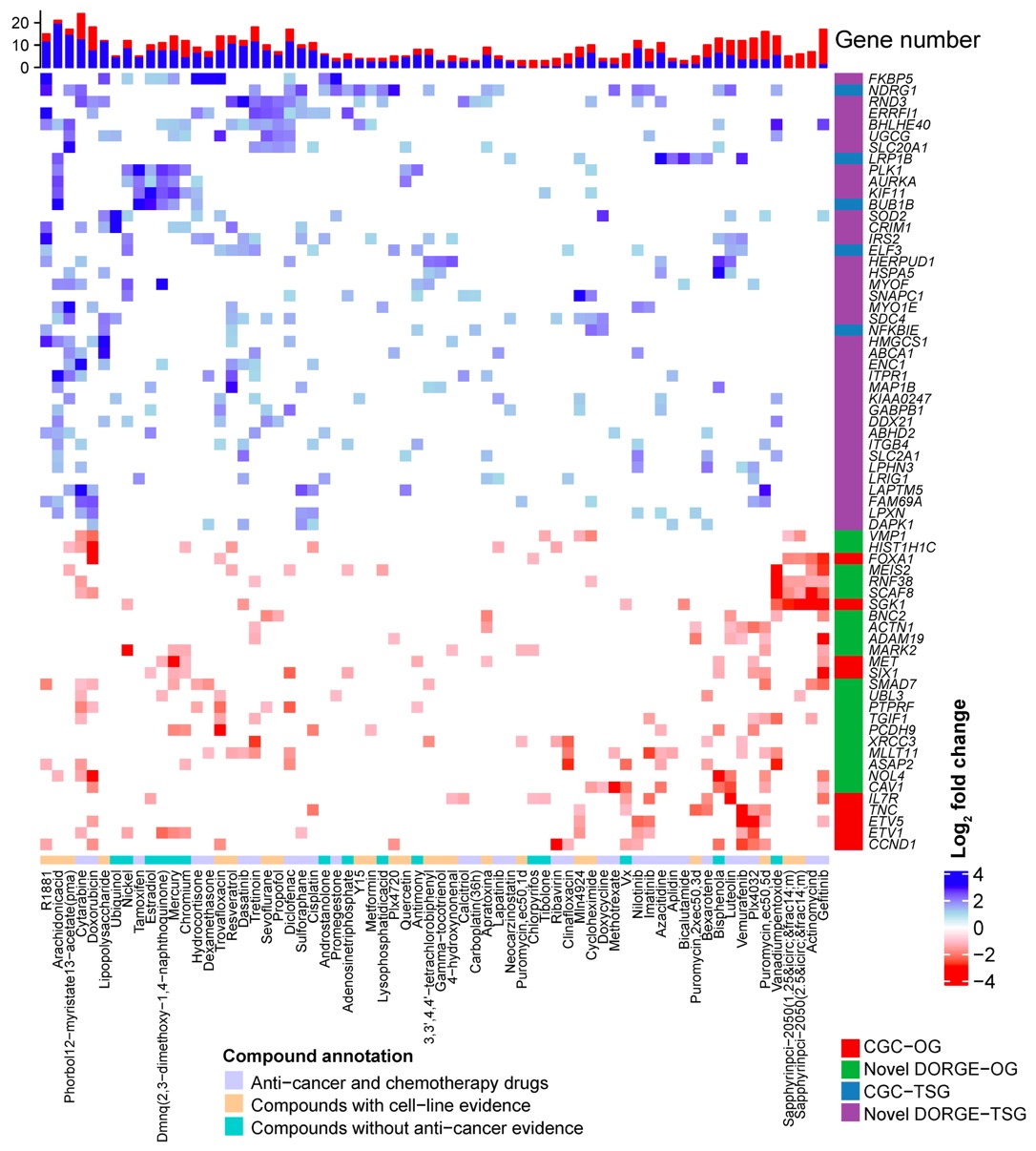

**Fig. S6. Heatmap of** **fold changes in expression for DORGE-predicted novel and CGC genes that are up- or down-regulated by drug perturbations.** Significant drug-gene relationships (*Q*-value < 0.05 and fold-change > 2) were obtained from the CRowd Extracted Expression of Differential Signatures (CREEDS) collection in the Drug Gene Budger (DGB) database. To reduce the number of genes/drugs in this figure, only those genes that are associated with at least seven compounds/drugs and compounds/drugs that are associated with at least three genes are presented. DORGE-predicted novel dual genes and CGC-dual genes were excluded from this analysis. The marginal stacked bar plot above the heatmap shows the total number of upregulated (blue) or downregulated (red) genes.

**Data file S1. Excel spreadsheet detailing the supplemental data including the candidate features, Spearman correlation table for features, histone modification data information, gene annotation, the DORGE predicted TSGs/OGs without occurring in cancer driver databases as well as the associated survival hazard ratios (HRs), and also the gene expression responses data for different drugs/compounds.**

**Data file S2. Excel spreadsheet detailing AUPRC values of eight classification algorithms under three class ratios, the feature groups, the DORGE prediction results and training gene annotations, evaluation of DORGE driver gene prediction on the OncoKB gene set, evaluation of DORGE TSG/OG prediction on the CGC gene list, the features used in different cancer driver prediction studies as well as the training feature profile.**
